# The pQBR mercury resistance plasmids: a model set of sympatric environmental mobile genetic elements

**DOI:** 10.64898/2026.03.27.714766

**Authors:** Victoria T. Orr, Ellie Harrison, Damian W. Rivett, Rosanna C. T. Wright, James P. J. Hall

## Abstract

Plasmids are extrachromosomal mobile genetic elements that can facilitate rapid bacterial adaptation by transferring genes between individuals. While plasmids are known to exist in diverse habitats and encode a range of traits, most of our knowledge about plasmids comes from clinically-associated antimicrobial resistance (AMR) plasmids that have already been recruited as vectors of drug resistance and have likely been shaped by strong selection for plasmid-encoded resistance. Here, we investigated 26 plasmids from the pQBR collection — a set of large, co-existing mercury resistance environmental plasmids isolated in *Pseudomonas* spp. from a field in Oxfordshire in the 1990s — and explored the ability of pQBR plasmids to mobilise novel chromosomally-encoded traits. New whole genome sequences for 25 plasmids confirmed that these soil-isolated plasmids are generally very large (140-588 kb), constitute at least five distinct genetic groups, and have relatives in various other *Pseudomonas* species and habitats. Despite significant nucleotide-level divergence, Groups I (pQBR103-like, ∼406 kb) and IV (pQBR57-like, ∼328 kb) showed remarkable ancient similarities in synteny and gene content both with one other, and with the PInc-2 family of plasmids known to mobilise clinically significant drug resistance in *Pseudomonas aeruginosa*. None of the pQBR plasmids sequenced to date harboured known AMR determinants, but putative phage defence systems and metal resistances were evident. Transposable elements, including the Tn5042 mercury resistance transposon, were responsible for significant structural variation within plasmid groups, consistent with a predominant role of transposons in rapidly remodelling plasmids. To experimentally test the ability of pQBR plasmids to spread new traits, we developed a novel transposon mobilisation assay which showed that certain Group IV pQBR plasmids were especially effective at acquiring the chromosomally-encoded transposon Tn6291, and that this mobilisation was likely due to specific plasmid factors rather than generic conjugation rate. Our work presents a tractable set of sequenced plasmids suitable for exploring the evolution and dynamics of gene acquisition by pre-AMR plasmids, and provides a key case study highlighting the pervasive interplay between plasmids and transposable elements that can drive microbial genome evolution.

**Repositories: github.com/jpjh/PQBR_PLASMIDS**

**Impact statement:** Plasmids can drive microbial evolution by acting as vectors for horizontal gene transfer. Because of their central role in disseminating antimicrobial resistance (AMR), plasmids are mainly explored as vehicles for AMR traits, meaning that our knowledge of the diversity and evolutionary dynamics of non-AMR plasmids is more limited. Here, we explore sequences from a set of mercury resistance plasmids isolated in *Pseudomonas* spp. from pristine agricultural land that lack AMR determinants. By providing new whole genome sequencing analyses we expand the set of sequenced pQBR plasmids to 26, finding globally dispersed relatives from clinical, environmental, and industrial settings, and identifying an ancient plasmid backbone shared amongst divergent modern environmental and clinical AMR plasmids. We experimentally verify the role of pQBR plasmids in readily mobilising chromosomal traits using a novel transposon mobilisation assay, which suggests that specific plasmid-transposon interactions may drive trait spread. Overall, our work expands our understanding of the role of environmental plasmids in mobilising and disseminating adaptive traits.

## Introduction

Plasmids are mobile genetic elements (MGEs) that exist separately from the chromosome as (usually) circular pieces of DNA, and can accelerate bacterial adaptation by transferring genes between individuals, predominantly via conjugation (1,2). Plasmid genomes can be conceptually divided into conserved ‘backbone’ sections that usually harbour genes directly responsible for plasmid fitness — such as those encoding machinery for plasmid replication and transmission — and a diverse and flexible accessory genome that can vary between otherwise closely-related plasmids and often encodes traits that affect the fitness of host bacteria, with indirect consequences for plasmid fitness (3). Accessory genes can provide a plethora of functions, including resistance to antibiotics, biocides, and metals; exotic metabolic and biodegradative pathways (4); and defence systems against other MGEs, including bacteriophage or even competing plasmids (5–7). Plasmid-mediated gene transfer poses a grave threat in the context of antimicrobial resistance in pathogens (8–10), but also offers opportunities for bioremediating polluted sites (11) or introducing novel traits into microbiomes (12), indicating the value of understanding the biological diversity of environmental conjugative plasmids.

Transposons (also known as transposable elements (TEs)) are regions of DNA encoding enzymes, called transposases, that interact with flanking DNA regions to catalyse the excision and re-integration of the transposon in a different site (2). Transposon activity can result in the proliferation of a transposon within a genome, and enable the transfer of the transposon into new genomes, such as in cases where the transposon inserts into another mobile genetic element, e.g. a conjugative plasmid. Here, the plasmid acts as a ‘vehicle’, enabling the transposon to escape the confines of a single bacterial lineage and mobilise into new backgrounds. Besides transposases, transposons can carry a cargo of accessory genes — in fact, many genes of interest within plasmid genomes, such as those conferring resistance traits, tend to be encoded on transposons (10,13). These interactions between plasmids and transposons can accelerate the spread of traits, as has been observed for resistance genes in hospital settings (14,15), because transposons enable genes to switch between plasmids, as well as on and off chromosomes (16). When comparing plasmid sequences across isolates, these patterns manifest as a conserved backbone region alongside ‘hotspot’ regions of great genetic diversity, as was observed for the IncP-1 plasmids (17).

Much of what we know about plasmids comes from antibiotic resistance plasmids isolated from clinical settings in the years following the widespread introduction of antibiotics in healthcare and agriculture. Strong selection for resistance can shape plasmid dynamics and evolution in various ways, for example, by driving high plasmid frequency in a population, selection can reduce opportunities for plasmid-mediated gene mobilisation (18) and favour mutations that increase copy number (19,20). Acquisition of resistance genes can also alter plasmid fitness effects (18,21). Comparative studies that span the pre- and post-antibiotic eras show that modern-day resistance plasmids have been recruited from a wider plasmid pool, and are closely related to historic plasmid backbones lacking resistance genes (22). Developing a broader understanding of plasmid dynamics and evolution therefore requires study of plasmids that have not undergone strong selection for specific resistance traits.

The pQBR collection is a set of sympatric plasmids isolated from the phytosphere of sugar beets grown in Oxford University Farm, Wytham, Oxford, UK (23–25). These plasmids were captured using exogenous isolation, whereby a rifampicin-resistant *Pseudomonas putida* UWC1 recipient (26) or kanamycin-resistant *Pseudomonas fluorescens* SBW25 recipient (27) was allowed to grow alongside the sugar beet microbiome before selection using rifampicin/kanamycin and mercuric chloride (HgCl_2_). The experimental design means that the original host of these plasmids in the sugar beet phytosphere is unknown, but at the moment of capture they were evidently mobilizable, able to replicate in *Pseudomonas* species, and confer phenotypic mercury resistance. An abundance of pQBR plasmids was captured this way — despite the source site having insufficient levels of mercury for positive selection and no known history of mercury pollution (24,25) — and were classified into groups based on restriction fragment length pattern (RFLP) similarities (23).

Experimental work performed on the pQBR plasmids over the last 30 years has significantly extended our knowledge of plasmid evolutionary ecology. This includes work on plasmid associations with plant roots (24) and biofilms (28), plasmid fitness (29), transfer (30), stability (31,32) and compensatory evolution (33–36). Sequences of pQBR103 (37), pQBR57, pQBR55 and pQBR44 (38), resolved in previous studies, revealed uncharacterised genetic novelty and hinted at vast unexplored diversity in the rhizosphere. There have also been studies focusing of on particular groups of genes, such as plant-inducible helicases in pQBR103 (39) and origins of replication in pQBR11 and pQBR55 (40,41). However, previous studies focused either on specific genes, or on individual divergent representatives of the restriction pattern groupings defined by Lilley *et al.* (1996). As a result, we lack a detailed understanding of diversity and evolution within and between more closely-related plasmids. Experimental studies have suggested the potential for rapid divergence between related pQBR plasmids by transposon acquisition and rearrangements (18), suggesting that these plasmids might be efficient vehicles for the mobilisation of chromosomally-encoded traits to other members of the soil microbiome.

Here, we increase the repertoire of sequenced pQBR plasmids, providing a snapshot of a community of co-existing plasmids in soil that have not been recruited for AMR gene carriage. We show that, consistent with earlier RFLP typing, the pQBR plasmids fall into distinct groups, with within-group structural variation driven by recent, prolific, transposon insertions. The backbones of each group include numerous genes and operons with predicted activities extending beyond the replication and conjugation of the plasmid, including chemotaxis, radical S-adenosyl-L-methionine (SAM) metabolism, and Type IV pilus formation. Using a novel assay, we show that the pQBR plasmids vary widely in their capacity for mobilising chromosomal transposons.

## Methods

### Bacterial culture

King’s B media (KB) was used to culture microorganisms. KB broth was made with 20 g Bacto-Proteose peptone No.3 (Difco), 1.5 g magnesium phosphate heptahydrate, 1.15 g potassium phosphate dibasic anhydrous, and 10 g glycerol (Honeywell) per litre. Agar (12 g/L) was added to make KB agar. All broth culturing, unless otherwise stated, was in 5 ml broth in 50 ml polypropylene tubes (Greiner Bio-One, 227261) within an Innova42 shaking incubator (New Brunswick Scientific) at 180 rpm and 28°C. Media supplementations with kanamycin were at 50 µg/ml, mercury chloride at 20 µM, streptomycin at 100 µg/ml and 5-bromo-4-chloro-3-indolyl-β-D-galactopyranoside (X-gal) at 50 µg/ml.

### Bacterial strains

Original pQBR plasmid carrying hosts were either *Pseudomonas putida* UWC1 (a derivative of *Pseudomonas putida* KT2440) or *Pseudomonas fluorescens* SBW25, but prior to this study all plasmids were transferred into *P. putida* UWC1 by conjugation as previously described (23,29). Streptomycin-resistant (SmR) *P. fluorescens* SBW25 with a *lacZ* marker was previously described by Hall *et al.* 2015 (38). *P. fluorescens* SBW25, harbouring a kanamycin resistance marker in Tn6291 (SBW25-Tn6291::KmR), was produced from a *P. fluorescens* SBW25 wild-type (27) by homologous recombination (42)(full details in Supplementary Methods). *Pseudomonas koreensis* P19E3 plasmid p1, referred to here as pP19E3.1 (43), was previously transferred into *P. fluorescens* SBW25 (10).

### Antibiotic resistance analysis

Antimicrobial resistance profiles were established by comparing plasmid free and plasmid carrying *P. putida* UWC1. Overnight cultures were spread (∼1 x 10^7^ cfu/ml) onto Mueller-Hinton agar (with and without 0.1 µg/ml HgCl_2_) using the M26/NCE multiple antimicrobial susceptibility ring (MAST Group Ltd.). Antimicrobials to which the *P. putida* UWC1 host was resistant to were removed from the analysis, along with Naladixic Acid to remove possibilities of spontaneous mutations, leaving colistin, kanamycin, streptomycin, and tetracycline as the test compounds. Zones of inhibition diameters were measured using digital callipers.

### Sequencing, assembly, and bioinformatic analysis of pQBR plasmids

Initial whole-genome sequencing of the pQBR plasmid collection (23,25) was performed on *P. putida* UWC1 pQBR strains using Illumina (2x250 bp, minimum 30x coverage, Illumina sequencing performed by MicrobesNG). To identify plasmid sequences, reads were first mapped against reference *Pseudomonas* chromosomes using ‘bwa-mem’ (44), and reads that did not map to the chromosome were extracted using SAMtools (45). This subset of reads was assembled using SPAdes v3.15.5 (46). Contigs that had long (>5kb) and deep coverage (>10x coverage) were identified as putative pQBR plasmid contigs and used for initial comparative analysis. These sequences were first compared with whole genome SPAdes assemblies in Bandage (47) to identify and resolve plasmid sequences. To attempt to close the remaining sequences, we used Oxford Nanopore Technology (ONT) sequencing. In some cases, ONT sequencing was performed on pools of DNA extracted from cells with different plasmids; combinations were selected so similar plasmids were not present in the same sample. DNA was isolated using the Masterpure^TM^ Complete DNA and RNA Purification Kit (Lucigen MC89010) and ONT bacterial genome sequencing was performed by Plasmidsaurus. ONT Flye (2.9.1-b1780) assemblies, polished with Medaka (1.8.0), were polished again using the corresponding Illumina reads with PolyPolish 0.5.0 (48,49), and direct repeats at the start and end of all assemblies were identified by ccfind 1.4.5 (50) and removed as assembly artefacts. These approaches produced closed plasmid sequences for 20/28 UWC1(pQBR) strains, with the remainder considered drafts; full details of the methods used to resolve all sequences provided in Supplementary Text.

All sequences were oriented using EMBOSS to place the first base of the first codon of a putative replicase at the first position on the forward strand. Replicase genes were identified by querying a BLAST database of putative replicases from previously-sequenced pQBR plasmids, with candidates chosen to facilitate visualisation of synteny where multiple candidates were available. Sequences were annotated using bakta (v1.8.2, full database v.5.0.0) (51) on metagenome mode. Example analysis scripts are provided on Github.

Heatmaps were produced in R (52–54) using mash distance (55) to cluster the sequences by similarity. Genome comparison plots were produced using tblastx and blastn, for intergroup and intragroup comparison respectively (56), and plotted using EasyFig (57). Sequences were uploaded to PHASTEST (v3.0) (58), TAFinder 2.0 (v3.0) (59), and DefenseFinder, including anti-defence finder (v2.2.0, Model 2.0.2) (60,61) and AMRFinderPlus (v4.0.23) (62), to identify potential phage insertions, putative toxin-antitoxin systems, putative genome defence and anti-defence systems, and predicted AMR genes respectively. Plasmid sequences were compared to the TnCentral database to identify any complete additional transposons present in the sequences (63). Relatives were identified by querying PLSDB (v2023_11_23_v2, sketch size 10,000) (64) and the complete genomes of PseudomonasDB (v22.1, sketch size 10,000) (65) using mash (55) with representatives from each group, and extracting sequences with an e-value < 1e-100. Candidates were subsequently filtered for matches with at least one blastn match >5 kb and e < 1e-40. Pangenomes were calculated using PIRATE (66) (v1.0.5) and nucleotide diversity for each gene cluster calculated using Mega (v11.0.13). Phylogenetic trees of plasmids and relatives were constructed in R using PIRATE core genome alignment, trimAL (v1.5.rev0) and IQtree3(v3.0.1)(52,67–71).

### Measuring and calculating conjugation and Tn6291 mobilisation rates

Plasmids were transferred into SBW25-Tn6291::KmR by conjugation. KB broth (5 ml) was inoculated with 50 µl of a 50:50 v/v mix of overnight cultures of a *P. putida* UWC1 plasmid donor strain and the recipient SBW25-Tn6291::KmR. Three separate mixes were established for each donor-recipient combination. Cultures were incubated overnight before samples were diluted and spread onto KB agar supplemented with mercury and kanamycin. Plates were assessed after 48 hours growth at 28°C. Single isolated transconjugants were re-streaked onto selective media, and isolated colonies were used to inoculate KB broth which was supplemented with glycerol (20% w/v) after overnight growth, and frozen at -80°C. Putative transconjugants were screened by PCR using plasmid-specific and recipient-specific primer pairs (Supplementary: Table S5). PCR was conducted with GoTaq Green G2 mastermix (Promega: M7822) using a 95°C denaturing temperature and annealing temperature of 58°C, both for 30 seconds and an extending temperature of 72°C for 1.5 minutes, with these steps repeating 30 times. A similar protocol was used to generate *P. fluorescens* SBW25-Tn6291::KmR(pP19E3.1). These strains gave an initial indication as to plasmid mobility and acted as donors in the subsequent assay to calculate transfer rates.

To measure plasmid conjugation and transposon mobilisation rates, single colonies of SBW25-Tn6291::KmR plasmid donor strains and *P. fluorescens* SBW25::SmR-*lacZ* recipient strains were separately cultured in 5 ml KB broth and incubated overnight. Each donor was mixed with a recipient at varying volumetric ratios to achieve approximate 1:1 start count ratios of donor and recipient. Three separate mixed cultures were made for each donor-recipient combination to produce three independent replicates, with two independent transconjugants tested for each plasmid. KB broth (5 ml) was then inoculated with 50 µl of the mixture. Start counts of donor and recipient were obtained by diluting and spreading samples of mixed culture onto KB agar containing X-gal. The *lacZ* gene enabled recipients to be distinguished from donors by production of a blue pigment on X-gal containing media. Cultures and plates were incubated overnight, after which the cultures were diluted and spread on selective and non-selective KB plates, also containing X-gal, to enumerate donors and recipients (non-selective plates), plasmid transconjugants (streptomycin and mercury supplementation), and transconjugants that had also received Tn6291::KmR (streptomycin and kanamycin supplementation). Single colonies of putative transconjugants, from the selective plates for both plasmid transfer and transposon mobilisation, were re-streaked on selective media and PCR (as previously described) was used to ascertain the presence of the Tn6291::KmR and plasmid.

To transfer the counts into rates, the following equation was used (72):

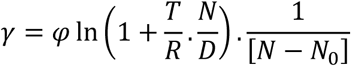

γ refers to the transfer rate (ml cell^-1^ h^-1^) (for either plasmid conjugation or transposon transfer), Ψ is the growth rate in (h^-1^); T, R, D are the number of transconjugants (or transposon mobilisations), recipients and donors respectively per ml. N is the final number of colony forming units per ml (cfu/ml) in the population and N_0_ is the starting number of cfu/ml in the population. The Simonsen equation assumes that donor, recipient, and transconjugant growth rates are equal, which is unlikely to be the case owing to known fitness costs of the pQBR plasmids (38). However, the approximate extended Simonsen method (ASM) (73), a more precise method, previously produced similar conjugation rates under our experimental conditions (36) and pilot studies indicated that our ASM protocol lacked the sensitivity to reliably detect transposon mobilisation events.

### Measuring plasmid fitness effects

Competitive fitness experiments were performed as described by Hall *et al.* 2015 (38), with details provided in Supplementary Methods, alongside details of growth curve analyses.

### Statistics

Data analysis was conducted in R (52) using the following packages: Tidyverse (53), forcats (74), multcomp (75), FSA (76). Figures were plotted using ggplot2 (77), patchwork (78), ggtext (79).

Ratio of transposon transfer compared to plasmid transfer was calculated by dividing γ for transposon transfer by γ for plasmid transfer, and log-transformed. If transposon mobilisation rates were lower than the limit of detection, their value was replaced by the limit threshold value before log-transformation. The effect on the ratio by plasmid and plasmid group, and effect of conjugation rate on transposon mobilisation rate were investigated using linear models, ANOVA and TukeyHSD post-hoc testing.

Comparative fitness values, w, were corrected by dividing by the mean of the control fitness values. Control values were not significantly different from 1 (mean w = 1.018; T-test, t=0.84, df=9, p=0.42). Corrected comparative fitness values were analysed using ANOVA with TukeyHSD post-hoc testing and plotted.

## Results

### Sequence analysis of pQBR plasmids reveals four groups of large plasmids capable of frequent, active mobilisation of transposable elements

The pQBR collection is a diverse group of mercury resistance plasmids captured from the same site in Oxfordshire by exogenous isolation over 3 years, using *Pseudomonas putida* UWC1 and *P. fluorescens* SBW25 recipients (23,25). Using whole-genome sequencing on 28 strains, including 24 with previously-unsequenced plasmids, we generated 20 complete plasmid assemblies and 5 draft assemblies. These were supplemented with the previously sequenced pQBR44 to give a total of 26 sequences for analysis (Supplementary: Table S1). The pQBR plasmids were originally classified into five groups (I-V) based on restriction fragment length polymorphism (23). At a global level (Fig.1), k-mer based comparison of the resolved plasmid sequences revealed that most of the plasmids (20/22) fell into these previously assigned groups (Supplementary: Table S1). With few exceptions, plasmids within each group had a high degree of identity and sequence similarity, whereas between groups there was little overall sequence similarity (Fig.1).

**Figure 1.**
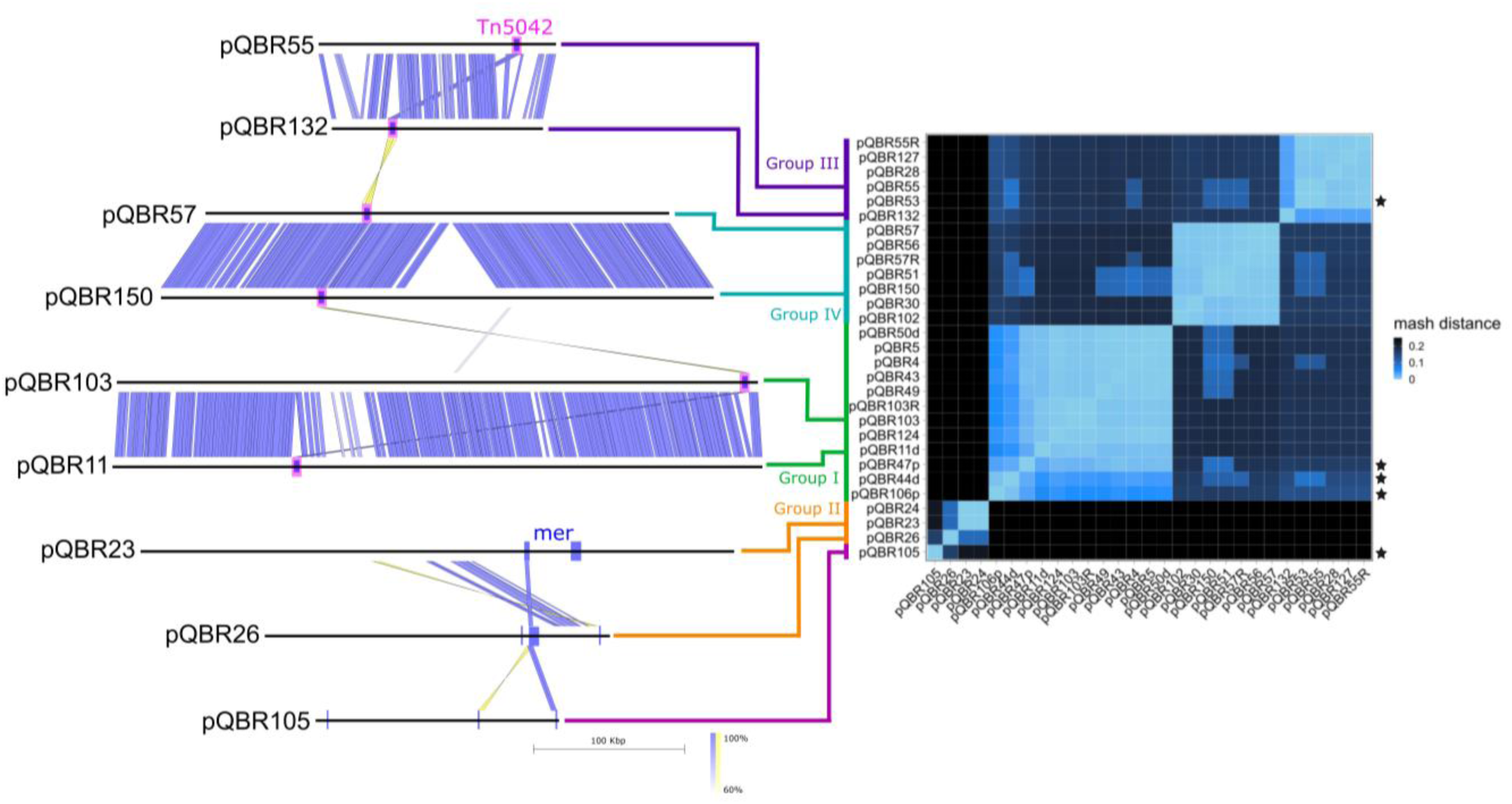
Sequence-based comparison of the pQBR plasmid collection shows distinct plasmid groups. On the right, a mash distance heatmap of all resolved plasmid sequences, with the brightness of each tile corresponding to the genetic similarity between each pair of plasmids. Stars indicate plasmids we were unable to conjugate into *P. fluorescens* SBW25 under our experimental conditions. On the left, a diagram showing representatives of each group indicates a high degree of conservation within each group, and a high degree of divergence between each group. Polygons indicate tblastx hits between adjacent sequences, with the intensity of the blue or yellow indicating sequence identity in the forward and reverse direction respectively (tblastx hits filtered for (>60% minimum identity, E < 1e-1000, 500 bp length of blast match). The location of the predominant mercury resistance transposon, Tn5042, is highlighted in pink, *mer* genes are indicated in blue.

Plasmid pQBR103, and by extension the Group I pQBR plasmids, were recently proposed to form a distinct group of *Pseudomonas* plasmids, termed PInc-17 or pQBR103-like (80). To investigate whether any of the other pQBR groups corresponded to the PIncs described by Nishimura *et al.* (80), we compared sequences to the canonical sequences of each of the PIncs. While the Group II plasmids produced significant matches — pQBR23 and pQBR24 matched the IncP_STY_ group (56% blastn coverage at >80% identity), and Group II plasmid pQBR26 matched the IncP-7 group (42% coverage) — there were no significant PInc matches for Group III (pQBR55-like) plasmids or Group IV (pQBR57-like) plasmids, which represent a novel grouping in the *Pseudomonas* plasmid PInc taxonomy.

The sequenced pQBR plasmids were generally large, and most are on the boundary of being megaplasmids (81), ranging in size from 140 kb (176 CDS, pQBR132) to 588 kb (712 CDS, pQBR49), and constituting 2.1% to 9.0% of the median genome size for *Pseudomonas* species (82) respectively (Fig. 2A). Furthermore, the pQBR plasmids are evidently mobile: most displayed conjugation between strains in laboratory settings in addition to the initial mobilisation into the exogenous isolation recipient (Supplementary: Table S1). All possessed a lower GC content compared to the average for *Pseudomonas* genomes (Wilcox test, p < 2.2e-16 for all plasmids; Fig.2B), thus they should not be considered immobile ‘chromids’ or sedentary secondary chromosomes.

**Figure 2.**
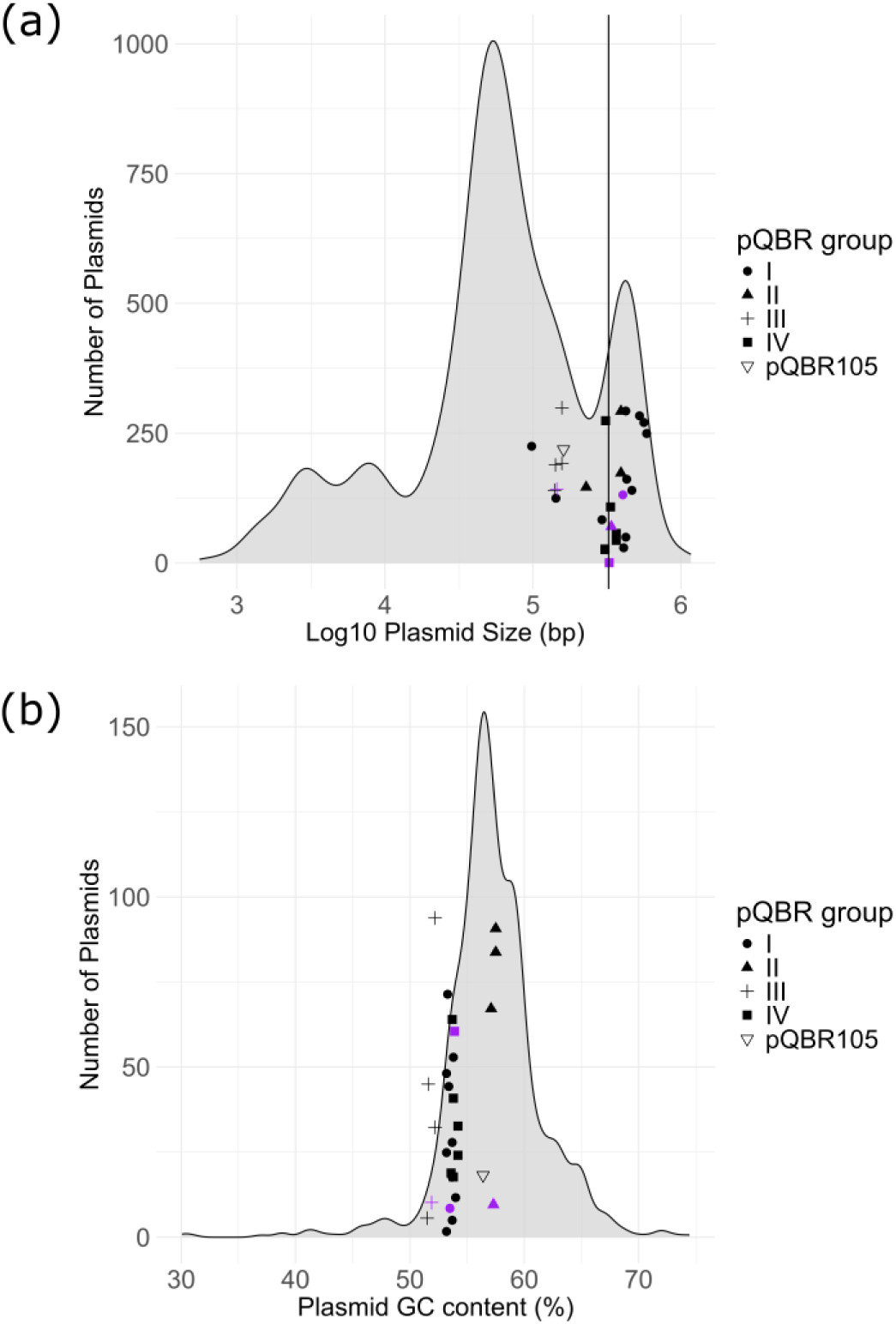
**A**. Plasmids within the pQBR collection are generally large. The size of each resolved pQBR plasmid is compared with a density plot of *Pseudomonas* plasmids in PLSDB (v.2024_05_31_v2). Vertical line represents 5% of median genome length of the complete genomes on PseudomonasDB. **B.** GC content (%) of pQBR plasmids compared to *Pseudomonas* plasmids within PLSDB. In both, figures plasmids from pQBR collection are represented by points jittered on the y-axis for visibility and group averages are indicated by purple points.

At the nucleotide level, few genes were conserved across plasmids from different groups. The remarkable exception was the presence of highly similar TEs found on plasmids from different groups. Most prominent was Tn5042, a 6,989-bp transposon with a *merRTPCAB* operon, previously identified on pQBR103, pQBR57 and pQBR55, and here found on all the other plasmids except Group II and pQBR105. These copies of Tn5042 all high sequence similarity and similar SNPs compared to the reference sequence (AJ563380.2) with few exceptions (see Supplementary Results). This strongly suggests that the transposon has moved relatively recently through the local plasmid population (23). In Group II, the mercury resistance transposon Tn5046 (10,118 bp) was identified in pQBR23 and pQBR24, sharing *merRTPCA* with ∼80% nucleotide similarity to Tn5042, and encoding *merDE* instead of *merB*. No intact mercury transposon with matches in TnCentral could be identified within pQBR26 or pQBR105, although unannotated mercury transposons are likely present (see Supplementary). The three pQBR plasmids that were lost before sequencing owing to chromosomal *mer* capture (pQBR1, pQBR8, and pQBR58, described in Supplementary Results) each left a transposon resembling the known *Pseudomonas* mercury resistance transposon Tn512 (83), which matched with >97% identity and >98% coverage, suggesting that this transposon might be more prone to chromosomal capture.

Three other full-length TEs were identified on plasmids across different groups: Tn4652 (Genbank accession AF151431.1, 17,029 bp), Tn6290 (BK010246.1, 42,031 bp), and Tn6291 (BK010245.1, 22,336 bp). These were generally well-conserved across the pQBR plasmids, although some contained minor divergences from their respective reference sequences (Supplementary: Table S2). These transposons are large, each with a cargo of genes with functions associated with metabolism, metal resistance, and transportation (see Supplementary Results). Transposons were sometimes present in multiple copies, most strikingly Tn6290 present in 3 copies in pQBR49. It is likely that these transposons were acquired during exogenous isolation as Tn4652 and Tn6290 are present in *P. putida* UWC1 chromosome while Tn6291 is present in *P. fluorescens* SBW25 (23,27). Of the 26 pQBR plasmids we analysed in the collection, 11 harboured isolation-associated TEs, demonstrating the readiness with which these MGEs can mobilise chromosomal elements and indicating the ability of pQBR plasmids to act as vehicles for horizontal gene transfer.

Using AMRFinderPlus, we did not identify any AMR genes within the pQBR collection and found no increased resistance conferred by any of the pQBR plasmids against colistin, kanamycin, streptomycin, or tetracycline in *P. putida* UWC1. The pQBR plasmids thus constitute a set of experimentally tractable, ecologically cohesive plasmids originating from a non-AMR, non-clinical context.

### Group I and IV pQBR plasmids have gene conservation despite extensive divergence, and are distant relatives of antimicrobial resistance plasmids

The Group I plasmids (n = 11, including the draft assemblies) were on average the largest plasmids in the pQBR collection, ranging between 98–588 kb, with a mean of 406 kb (Supplementary: Table S1). As previously observed for pQBR44 (38), plasmids pQBR47 and pQBR106 are likely to represent truncated variants: these three plasmids were substantially smaller than the others in the group, lack several of the key functional regions predicted for other Group I plasmids (including conjugative transfer) and were not self-mobilisable in our experimental assays (Supplementary: Table S1). The remaining Group I plasmids shared extensive identity and synteny, with most structural variation due to TEs (Fig. 3A). Mercury transposon Tn5042 was located in a syntenic region in all Group I plasmids, except for pQBR103 and pQBR11 in which the locations were unique. Plasmids pQBR4, pQBR5, pQBR43, pQBR47, pQBR49 and pQBR50 carried transposon Tn6290, sometimes with multiple copies. Most of these were in different locations, but pQBR4, pQBR47, pQBR49, and pQBR50 had at least one conserved insertion site (Fig.3A).

**Figure 3.**
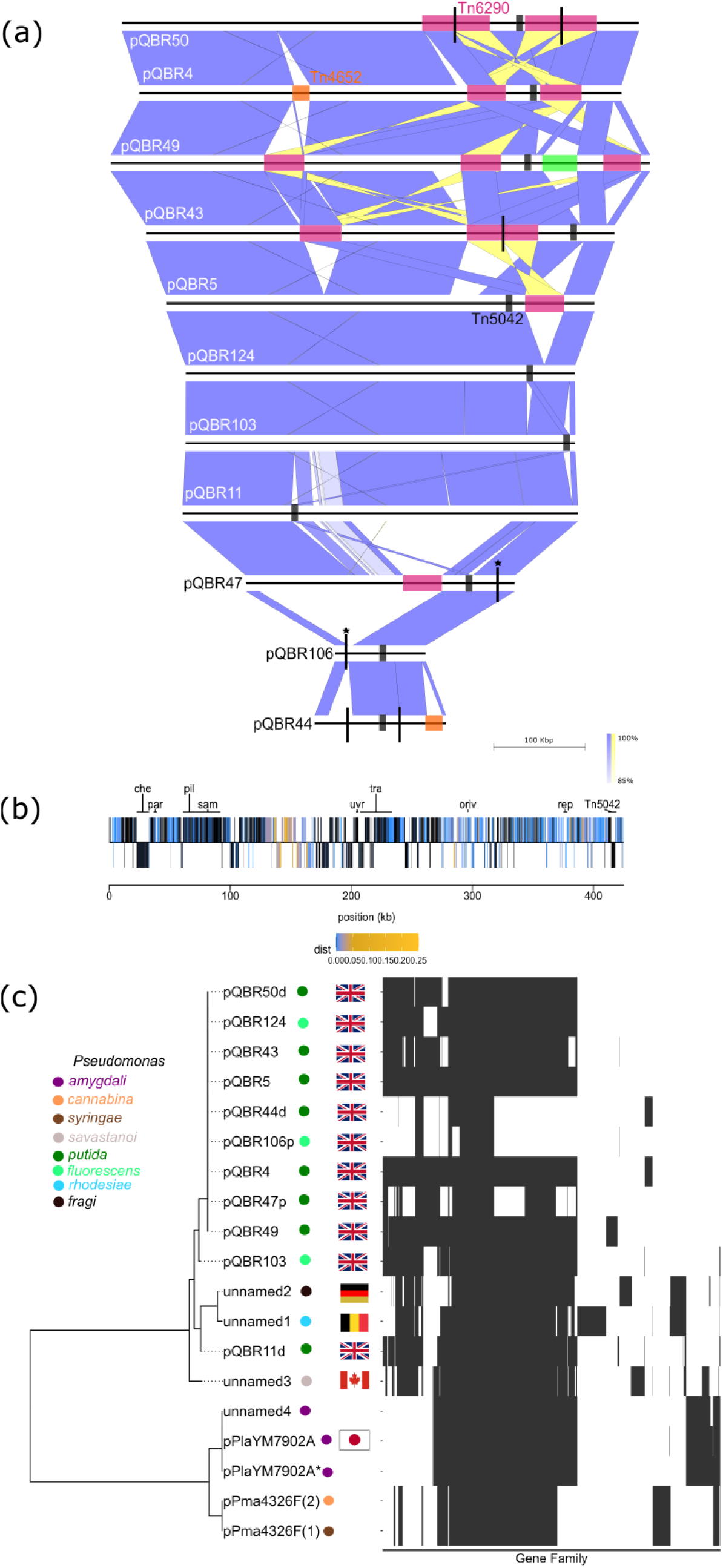
Group I pQBR plasmids share a conserved backbone but have experienced high levels of transposon activity alongside truncations. **A.** blastn comparison of Group I pQBR plasmids, with transposons highlighted in pink (Tn6290), orange (Tn4652) and black (Tn5042). The green highlighted region in pQBR49 is the unique region containing Erebus defence system. Lines extending above and below lines on these sequences indicate location of contig breaks, due to Tn6290 presence interfering with assembly, and lines with a star above indicate breaks due to potential chromosomal integration. **B.** Comparison of gene conservation amongst Group I plasmids. Raw nucleotide divergence (p-distance) was calculated across homologues for each pQBR103 gene. Genes with higher divergence across the group are shown in more intense shades of yellow and blue, with invariant genes coloured in black. Genes above or below the line represent genes on the forward or reverse strand. **C.** Phylogenetic tree of the core genome and heatmaps showing gene presence/absence across the Group I plasmids and relatives from PLSDB and PseudomonasDB. Note that pPma4326F(1) and pPma4326F(2) are from different host strains, *P. syringae pv. maculicola* (NZ_CP047261.1) and *P. cannabina pv. alisalensis* (NZ_CP084324.1) respectively.

Examining neighbouring genes to each Tn5042 copy revealed that TE acquisition likely disrupted plasmid genes: a SmpA-OmlA domain-containing protein, which was intact in pQBR11 (PQBR11d_02450) and pQBR103 (PQBR103_02355), was present only in truncated or pseudogenized form due to the insertion site shared by pQBR4 and relatives. TE insertion appeared to have disrupted a Transglut-core3 domain-containing protein in pQBR11 (PQBR11d_00620), and in pQBR103, a DUF913 domain-containing protein (PQBR103_02645). An additional orphan ParB protein was noted in pQBR103 and pQBR11, and pQBR49 contained a unique region, encoding putative phage defence system Erebus (Fig. 3A) (84). The region unique to pQBR49 also contains several phage/MGE related genes, including a holin (PQBR49_03095), a phage protein (PQBR49_03025), four Tyr recombinases (PQBR49_03030, PQBR49_03045, PQBR49_03055, PQBR49_03060) and a transposase (PQBR49_02960), suggesting this region is a degraded remnant of another MGE (no intact phages were detected on any plasmids). MOBsuite could not identify the mobility or relaxase genes within Group I, or match them to known MOB gene families, but most Group I plasmids were experimentally conjugated into *P. fluorescens* SBW25 successfully (Supplementary: Table S1).

Using the backbone sequence of pQBR103 as a query, we used mash (55) to search public databases for related plasmids. We found 13 complete plasmid sequences with an average 85-93% nucleotide identity to pQBR103, with a 300-400 kb shared backbone in non-truncated variants. All sequences were associated with *Pseudomonas spp.* and most originated from plant-associated environments or contaminated soils, except for two: a *P. rhodesiae* plasmid and a *Pseudomonas fragi* plasmid isolated from a brewery (85) and raw milk respectively (86) (Supplementary: Table S3).

The Group IV plasmids (n = 6) ranged between 307–366 kb (328 kb average) and were all conjugative (Supplementary: Table S1). The six plasmids showed extensive identity and synteny, alongside TE activity. Tn5042 was present in syntenic positions in pQBR51, pQBR56, pQBR57 and pQBR150, potentially disrupting an efflux RND transporter periplasmic adaptor subunit and flanked by DNA helicase and a transcriptional regulator. A Tn5042 insertion site was shared between pQBR30 and pQBR102, with no obvious gene disruption when comparing with a syntenic site on the other plasmids. Two plasmids (pQBR51, pQBR150) contained Tn4652 and Tn6290 in the same location and tandem arrangement, whereby Tn4652 appeared to have inserted within Tn6290 at base 40,829 in the transposon reference sequence to generate a nested structure. Tn6291 was found only in pQBR102, flanked by a DNA helicase. Beyond TE activity, minor variation in gene content mainly came from pQBR30 and pQBR102 (details in Supplementary Results).

As with the Group I plasmids, we attempted to identify related plasmids for Group IV, and found only a draft *P. putida* genome of unknown origin (NZ_JANHLM010000005.1). MOBsuite was unable to predict mobility correctly for Group IV and relaxases could not be matched to known MOB gene families.

The Group I and Group IV plasmids each harbour a conserved backbone ∼200 kb in length. We investigated sequence variation across the conserved genes (Figures 3B and 4B). The most divergent genes across Group I were mainly genes of unknown function, several of which encoded putatively membrane-associated proteins. Within Group IV, divergent genes included a transcriptional regulator, a serine/threonine protein kinase, a putative bacteriophage protein, a YrdC-like domain-containing protein, and two leader peptidases. These may represent more rapidly-evolving loci, perhaps involved in coevolutionary interactions, but the unknown function of these genes makes further speculation difficult.

**Figure 4.**
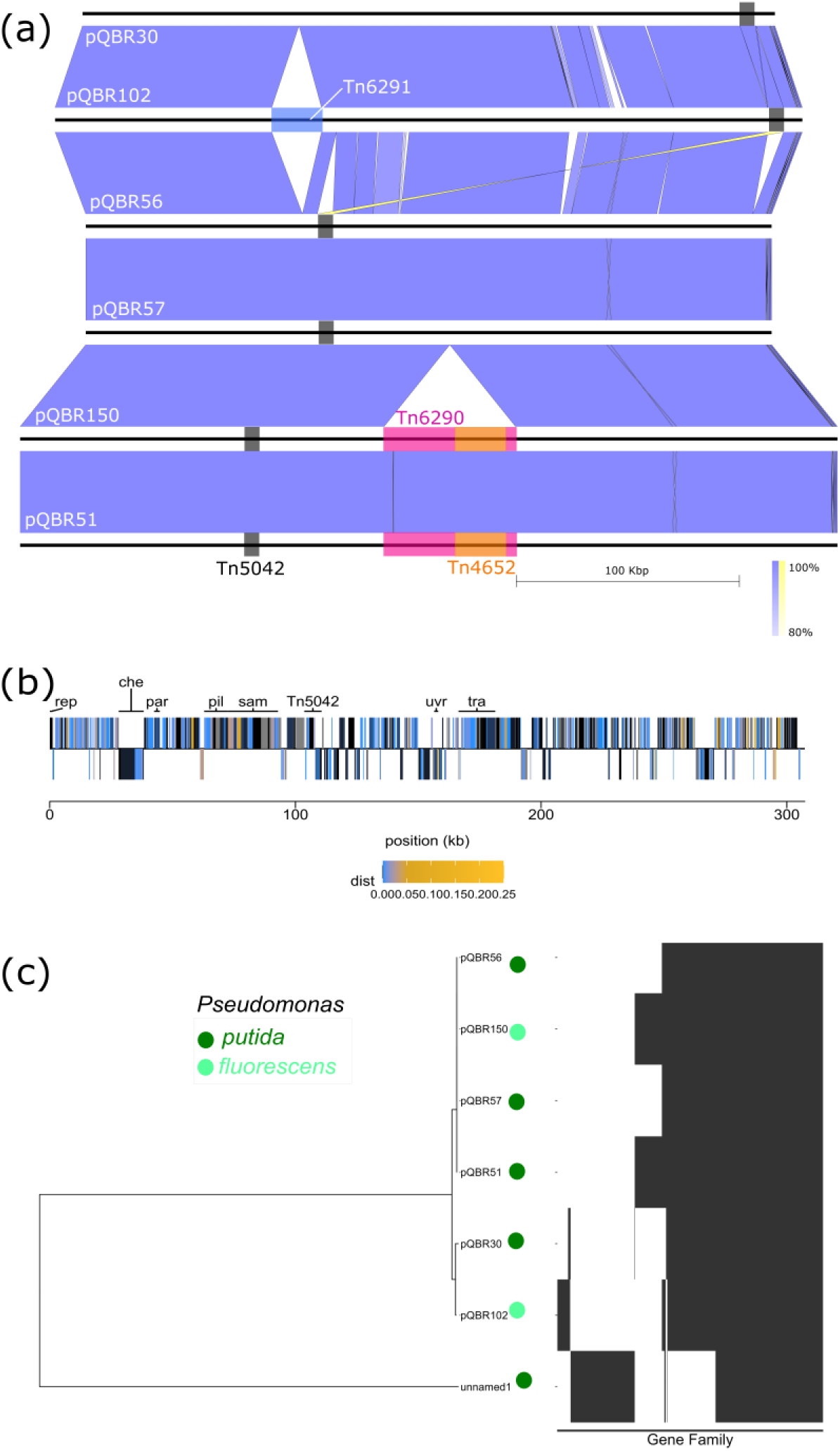
Group IV pQBR plasmids shared a conserved backbone. **A.** blastn comparison of Group IV pQBR plasmids, with transposons highlighted in blue (Tn6291), pink (Tn6290), orange (Tn4652)and black (Tn5042). **B.** Comparison of gene conservation amongst Group IV plasmids. Raw nucleotide divergence (p-distance) was calculated across homologues for each pQBR57 gene. Genes with higher divergence across the group are shown in more intense shades of yellow and blue, with invariant genes coloured in black. **C.** Phylogenetic tree of the core genome and heatmaps showing gene presence/absence across the Group IV plasmids and the relative identified in PseudomonasDB.

Though divergent at the nucleotide level, the Group I and Group IV backbones contain many similar genes arranged in similar order, encoding putative functional pathways such as a chemotaxis phosphorelay system, a type IV pilus, and radical SAM-metabolising enzymatic clusters (38). Divergent, yet similar, backbone regions were also found to be present in the IncP-2 plasmids, including the multi-drug resistance plasmid pBT2436 and the environmental megaplasmid p1 (from strain *P. koreensis* P19E3) (10), and the *P. shirazica* plasmid pJBCL41 (87). We compared the backbone regions of Group I, Group IV, IncP-2, and pJBCL41-like plasmids and whilst highly divergent at the nucleotide level (43-50% nucleotide identity), tblastx comparisons clearly highlighted clusters of 53 kb and 33 kb of homologous genes with similar predicted functions (Fig.5). Together, our results show an ancient plasmid backbone shared amongst environmental and clinical plasmids worldwide, capable of readily mobilising traits including antibiotic and metal resistance.

**Figure 5.**
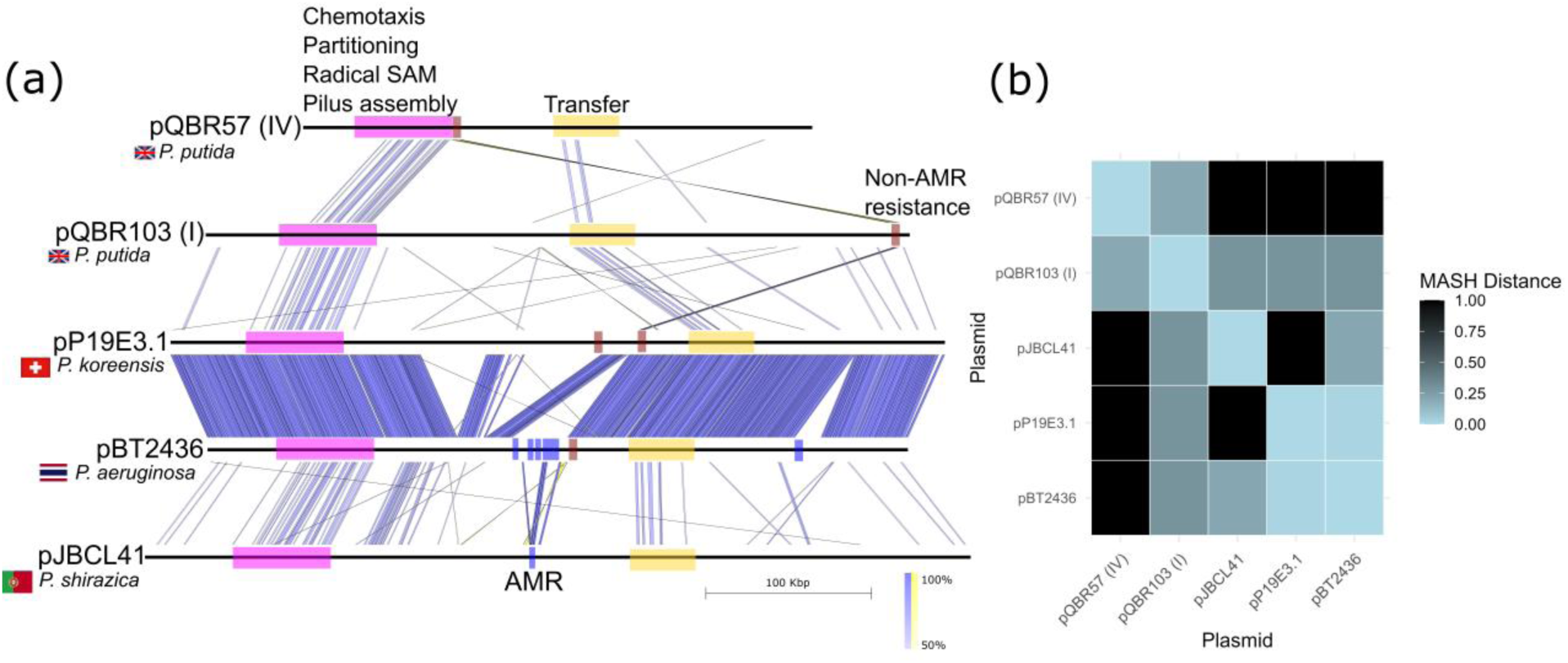
Group I and Group IV plasmids have divergent, globally-dispersed relatives. A comparison of plasmids pQBR103 (Group I), pQBR57 (Group IV), p1 from *P. koreensis* P19E3 and pBT2436 from *P. aeruginosa* BT2436 (IncP-2) and pJBCL41 from *P. shirazica* FFUP_PS_41. **A.** tblastx comparison of plasmids (>50% minimum identity, E < 1e-1000, minimum blast match length 200 bp) with shared regions of interest highlighted. Antibiotic resistance (AMR) and non-AMR resistance genes, as predicted by AMRFinderPlus, are highlighted in dark blue and red respectively. **B.** Heatmap of sequence similarity based on Mash distance.

### Sequence comparisons of other pQBR plasmid groups

In Group III (n = 4), plasmids ranged from 140–157 kb (mean 145 kb). Again, we identified TE activity (from Tn5042 and Tn4652), and, in contrast to Groups I and IV, significant divergence in the backbone sequences, particularly with pQBR132, which had lost various genes from across the plasmid and showed lower overall sequence identity (Fig. 6A). Of note, pQBR132 did not encode a MazEF toxin-antitoxin system shared amongst other Group III plasmids (Supplementary: Table S1), while the genes for paREP8 (potentially involved in cell defence (88)), and a dystrophin (pQBR132_00470) were apparently truncated. Tn5042 was present in a syntenic location within pQBR53, pQBR55 and pQBR127, except for a DNA binding protein in pQBR127 which may have been disrupted by the transposon (PQBR127_00665). The other two Group III plasmids have different insertion site locations. Plasmid pQBR53 in Group III has transposon Tn4652 inserted into a putative *traC* CDS and pilus assembly protein which may have impacted its ability to conjugate, as it was the only Group III plasmid we were unable to conjugate into *P. fluorescens* SBW25 (Supplementary: Table S1). The most divergent genes across the Group III plasmids were related to DNA processing, transcriptional regulation, and metabolism (Fig. 6B).

**Figure 6.**
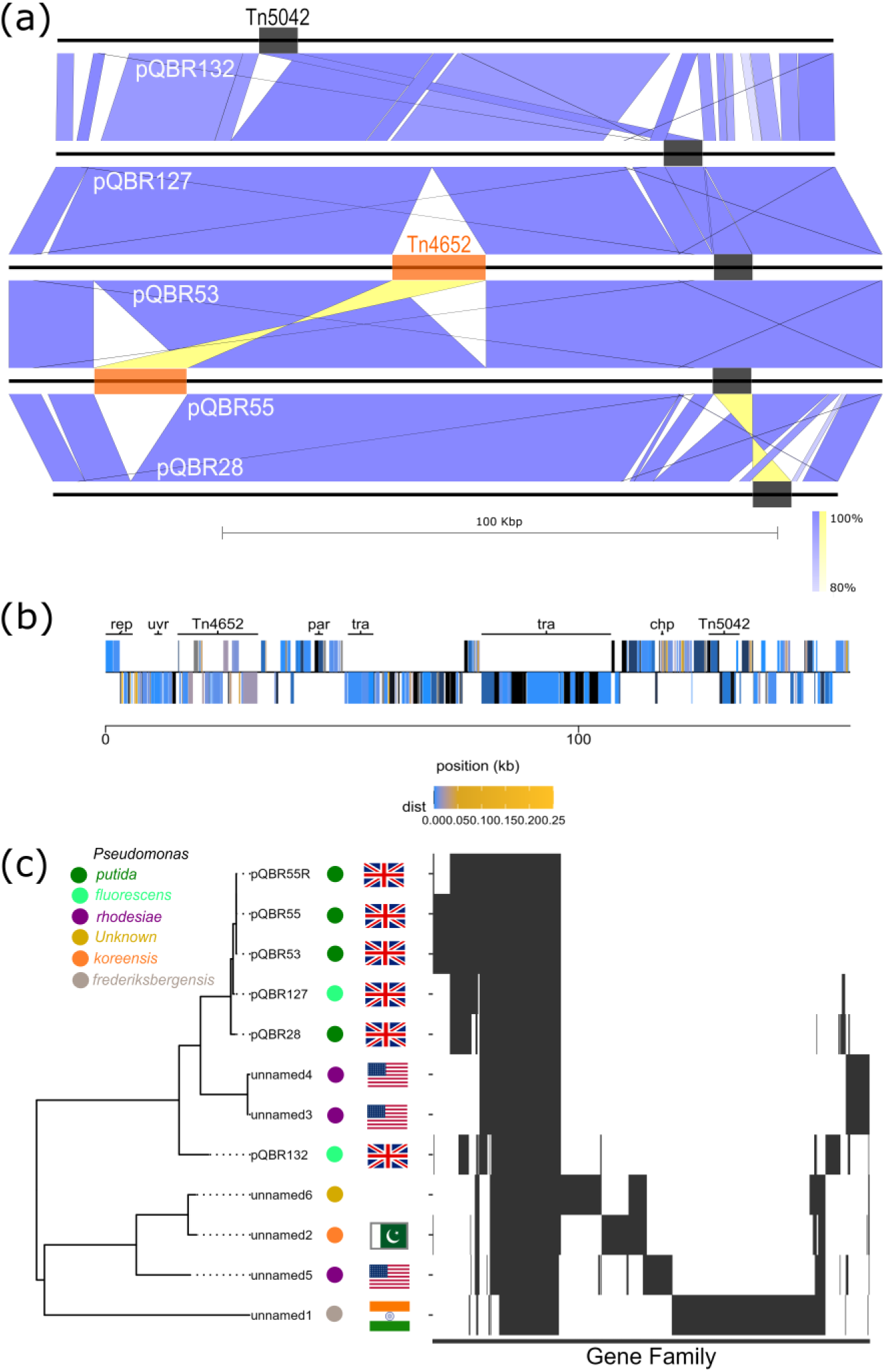
Group III pQBR plasmids show more backbone divergence than Groups I and IV. **A.** blastn comparison of Group III pQBR plasmids, with transposons highlighted in blue (Tn6291), pink (Tn6290), orange (Tn4652) and black (Tn5042). **B.** Raw nucleotide divergence (p-distance) was calculated across homologues for each pQBR57 gene. Genes with higher divergence across the group are shown in more intense shades of yellow and blue, with invariant genes coloured in black. **C.** Phylogenetic tree of the core genome and heatmaps showing gene presence/absence across the Group I plasmids and relatives from PLSDB, and PseudomonasDB.

We identified six unnamed plasmids in public databases that matched Group III, mainly matching with putative replication and transfer loci (Supplemental Results). Using tblastx, these related plasmids shared core function genes, including partitioning, conjugal transfer, regulatory and DNA replication genes. The two plasmids of known origin, from *P. koreensis* and *P. frederiksbergensis,* were isolated from soil and a glacial stream respectively (Supplementary: Table S4). *P. resinovorans* plasmid pCAR1 (AB088420), previously identified to share some matching regions with pQBR55 (38), had greater similarity to the Group II plasmid pQBR26, with all three encoding MOBH relaxases.

Two of the Group II (n = 3) plasmids (pQBR23 and pQBR24, Average Nucleotide Identity (ANI > 99.9%) were highly similar, with the other (pQBR26) more divergent (ANI ∼94.06) (89). All three contained multiple toxin-antitoxin systems and defence systems (Supplementary: Table S1). All three plasmids were predicted to be conjugative by MOBsuite, with pQBR23 and pQBR24 encoding MOBF and MOBP relaxases and pQBR26 encoding a MOBH relaxase. MOBsuite also predicted that *Pseudomonas. taiwanensis* VLB120 pSTY (isolated from forest soil, Germany) was related to pQBR23 and pQBR24; and pQBR26 was related to carbapenemase-encoding *Pseudomonas aeruginosa* plasmid p1160-VIM (China) (Fig.7 A&B). In pQBR26, a CRISPR-Cas system (subtype IV-A) and anti-CRISPR system were identified by DefenseFinder (Supplementary: Table S1). Spacer sequences from the CRISPR-Cas in pQBR26 revealed matches to genomes of plasmids in *P. aeruginosa* and *P. paraeruginosa* (CP173144.1, CP029091.1, CP027167.1, 128–166 kb in size) and one *P. rhodesiae* chromosome (LT629801.1). Comparison of these plasmids to representative from the pQBR groups (pQBR57, pQBR103, pQBR55, pQBR23, pQBR26 and pQBR105) showed minimal nucleotide similarity, suggesting that the pQBR26-encoded CRISPR-Cas is involved in targeting dissimilar plasmid competitors.

**Figure 7.**
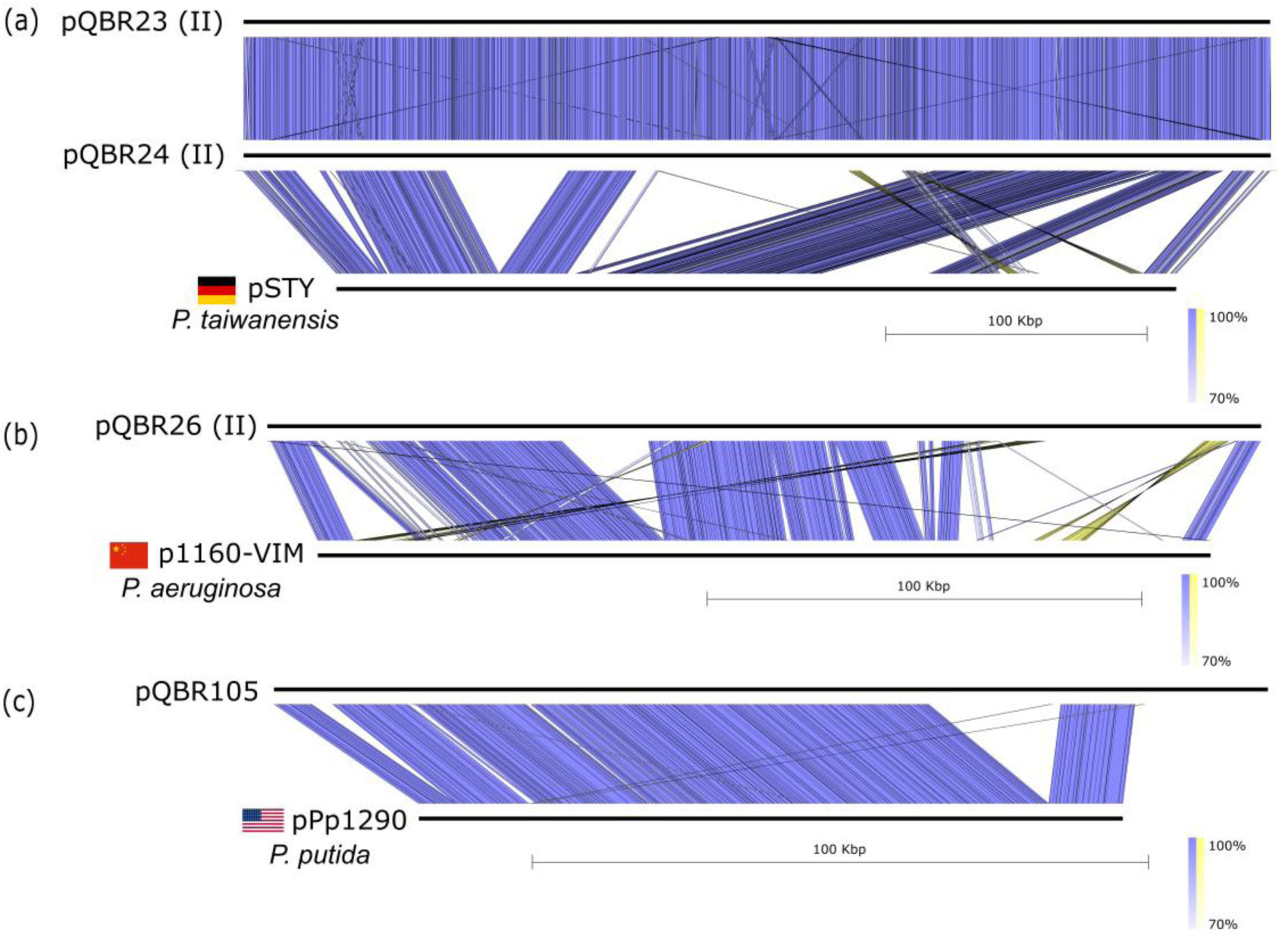
Group II and pQBR105-like plasmids have globally-dispersed relatives identified by mob_typer. **A**. tblastx comparison of Group II plasmids pQBR23 and pQBR24 with *P. taiwanensis* VLB120 pSTY. **B.** Group II plasmid pQBR26 and *P. aeruginosa* p1160-VIM. **C.** Plasmid pQBR105 and *P. putida* pPp1290. All tblastx comparisons were filtered >70% minimum identity, E < 1e-1000, minimum blast match length 100 bp.

Plasmid pQBR105 did not share enough relatedness to any of the other pQBR plasmids to be classified into Group I-IV. Although containing mercury resistance genes and many transposases, we were unable to identify any known transposons registered with TnCentral. plasmid pQBR105 is predicted to be conjugative by MOBsuite, with MOBP relaxases identified and relatedness to *Pseudomonas putida* pPp1290 isolated from an orchard leaf in California, USA (Fig.7C), but we were not able to mobilise it to *P. fluorescens* SBW25 in our assays.

### Plasmid mobilisation of chromosomal traits varies across pQBR plasmids, and is not determined by conjugation rate

Previously, pQBR57 was observed to be an effective vehicle for inter-species chromosomal transposon mobilisation (18). The presence of diverse TEs across different pQBR plasmid groups indicates that these plasmids may be effective vehicles for mobilising and disseminating TE-borne traits. To investigate the capacity of different pQBR plasmids to mobilise a chromosomally-encoded TE, we tracked the mobilisation of Tn6291 from the *P. fluorescens* SBW25 chromosome to a naive host. Using differential selective markers, we were able to distinguish plasmid conjugation from plasmid-mediated transposon mobilisation. Alongside the pQBR plasmids we included the pP19E3.1 (43), which shares distant backbone similarity to pQBR57 and pQBR103 and has close relatives implicated in the global spread of antibiotic resistance, as described above.

We hypothesised that transposon mobilisation would vary with conjugation rate, with the most conjugative plasmids being most effective at mobilising Tn6291. However, we did not find evidence of a significant relationship between conjugation rate and transposon mobilisation rate (Figure 8: F_(1,69)_=1.173, p=0.28), with ratios of plasmid conjugation rate to transposon mobilisation rate varying between plasmids and plasmid groups by several orders of magnitude (Supplementary: Figure S1, F_(8,62)_=34.88, p<0.01 for plasmids; F_(4,66)_=28.18, p<0.01 for Groups). For example, pQBR132 (Group III) had a strikingly lower ratio compared with the other plasmids (p<0.01), being ineffective at mobilising Tn6291 despite a relatively high conjugation rate, while pQBR102 and pQBR57 (both Group IV) were generally most effective at mobilising Tn6291. Of every ∼2359 pQBR102 transconjugant colonies and ∼7255 pQBR57 transconjugant colonies, one had also acquired Tn6291::KmR. To investigate whether mobilisation rate correlated with plasmid costs, we performed growth curves and competitive fitness assays, but did not identify any correlation between plasmid fitness effects and the efficiency of the plasmid in mobilising a chromosomal locus, suggesting that the striking differences in transposon acquisition and mobilisation are due to more specific differences in plasmid activity or gene content (Supplementary: Figure S2).

**Figure 8.**
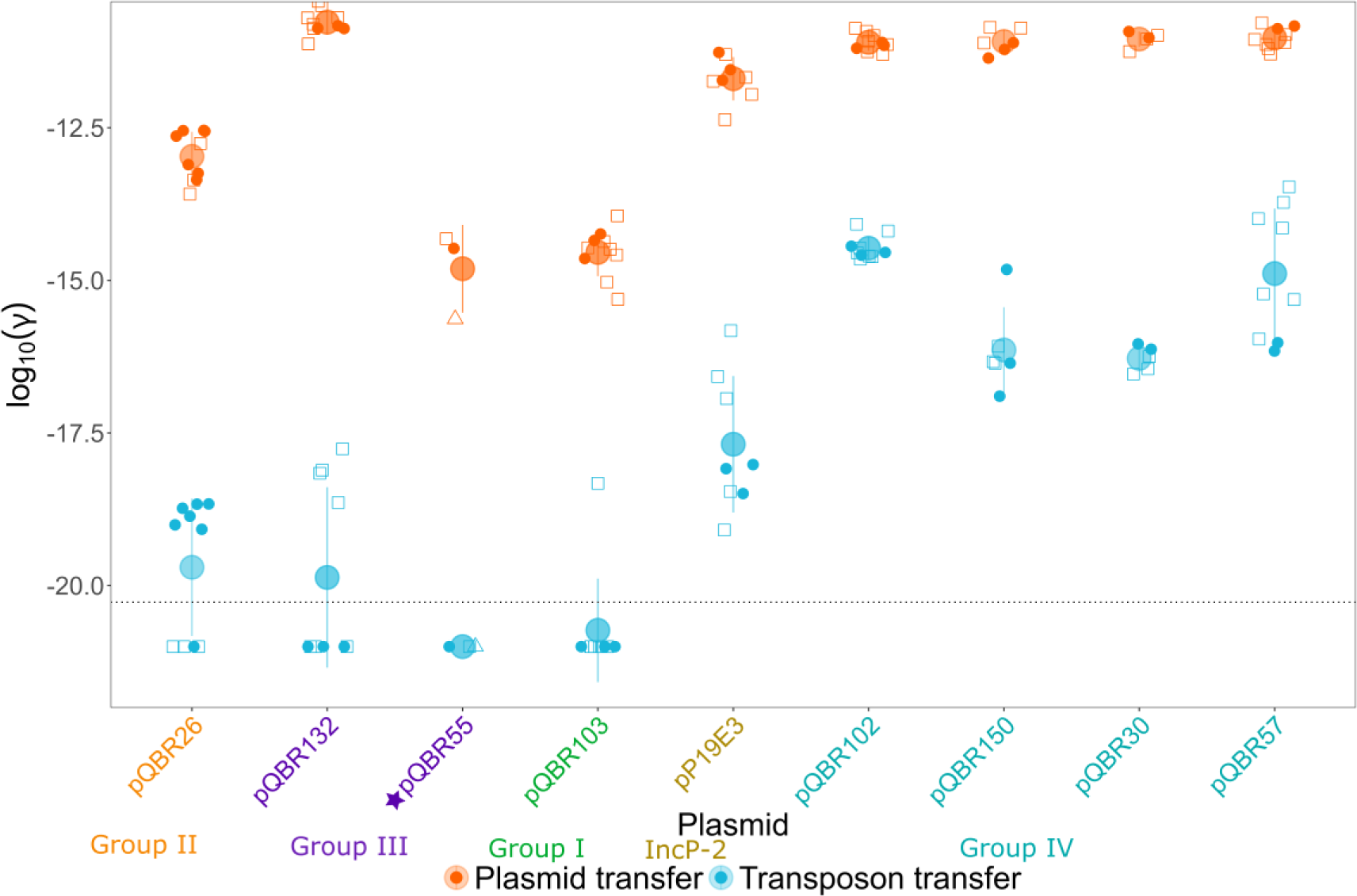
Conjugation rate is not a good predictor of transposon mobilisation. The lower plot shows transposon transfer rates (blue) and plasmid transfer rate (orange). Dotted line represents estimated limit of detection, calculated using average growth rate and if a single transconjugant colony was counted. Large circular datapoints represent mean of log10 transformed transfer rate values for that plasmid and bars represent the standard deviation. This figure represents a summary of two experiments performed a month apart, with 2-3 different donor transconjugant strains for each plasmid. Colours of plasmid labels represents the group the plasmid belongs to, labelled beneath. Star (pQBR55) indicates that data for this plasmid was collected separately from the rest of the dataset under similar experimental conditions.

## Discussion

The pQBR plasmids were isolated from a field growing sugar beets in Oxfordshire, UK (23,24,29) in the 1990s and represents a collection of environmental, co-occurring plasmids that do not encode antibiotic resistance genes. This collection therefore represents a valuable system for understanding the natural ecology of ‘wild’ plasmids from non-clinical settings. Here, we have analysed 26 pQBR plasmids, including 22 novel sequences, providing a detailed insight into plasmid genetics alongside conjugation and gene mobilisation phenotypes, illuminating the diversity that can be found both within a plasmid community and between closely-related plasmids, as well as the rapid plasmid evolution that can occur through the activity of TEs. The pQBR plasmids were isolated from a field growing sugar beets in Oxfordshire, UK (23,24,29) in the 1990s and represent a collection of environmental, co-occurring plasmids that do not encode antibiotic resistance genes. This collection therefore represents a valuable system for understanding the natural ecology of ‘wild’ plasmids from non-clinical settings. Here, we have analysed 26 pQBR plasmids, including 22 newly-sequenced plasmids, providing a detailed insight into plasmid genetics alongside conjugation and gene mobilisation phenotypes, illuminating the diversity that can be found both within a plasmid community and between closely-related plasmids, as well as the rapid plasmid evolution that can occur through the activity of TEs.

The pQBR plasmids are relatively large, consistent with other plasmids from soil (90). Genome-based clustering satisfyingly recapitulated prior RFLP-based analyses (23) to place the sequenced plasmids into distinct groups. Superficially, these analyses suggested that pQBR plasmids of different groups have little in common beyond their ability to confer mercury resistance. However, closer analyses found distant resemblances when comparing plasmids of Group I and IV despite low nucleotide identities, and similarly, both of these groups had distant similarities with both the clinically-important PInc-2/IncP-2 resistance plasmids and the antimicrobial resistance megaplasmid pJBCL41. The presence of similar genes in a similar order — but highly diverged at the nucleotide level — speaks to ancient conservation of function across these distant plasmid taxonomic units, which are found in environments including the infected lung (10,91), industrial processing (92,93), and soil habitats, and across diverse species of *Pseudomonas* (Fig.3A, Fig.5 & Supplementary: Table S3). Interestingly, these ancient conserved genes seem to encode features that are not obviously directly related to plasmid fitness, including chemotaxis, adherence and motility (Type IV pilus), and arylsulfatase/radical SAM/iron-sulfur cluster metabolism. While the functional role of these genes is yet to be determined experimentally, this observation highlights that plasmids likely have long-standing and nuanced life histories that extend beyond an existence as either simple vectors of functional traits, or entirely selfish parasitic agents.

A previous comparison of plasmids from the pre- and post-antibiotic era showed that 77% of pre-antibiotic era plasmids continued to circulate and maintain a recognisable backbone in modern times, in many cases having acquired one or more resistance genes (22). We did not identify any known AMR genes on any of the pQBR plasmids, and plasmid carriage did not affect any antibiotic resistance phenotypes tested. The pQBR collection therefore represents a tractable set of modern environmental plasmids that have not yet been recruited as vehicles for antibiotic resistance, opening the opportunity to explore the biology of pre-resistance plasmids, and experimentally study the dynamics of AMR gene acquisition by plasmids.

Within each group, significant structural differences emerged from the variable presence of TEs. Plasmids were selected using mercury, and it is likely that, thanks to TE activity, mercury resistance spread through this local plasmid community, effectively labelling a cross-section for retrieval using exogenous isolation, and providing another example of the predominant role that transposon activity can play in rapid plasmid evolution. In many cases, TEs were likely acquired during the process of exogenous isolation, as transposons identified were often present in the isolation host strains and dramatic abiotic changes associated with exogenous isolation may have triggered transposition. For example, presence of metals within media has been shown to enhance the activity of Tn4652 (94). However, there was variation in the patterns of different transposons between groups, potentially hinting at barriers or limitations emerging from plasmid host range or genetic content. For example, Tn6290 was prolific in its insertion within Group I, with 45% (6/11) of the sequences containing this transposon, often with multiple copies. Whilst, in Group IV, Tn6290 was found in 33% (2/6) of the plasmids with a single copy, a nested variant also harbouring Tn4652. The presence of many TEs and transposases in these sequences, and their apparently recent and repeated acquisition, allude to the readiness of these plasmids to acquire traits through transposition.

TE acquisition may not always be beneficial, as insertion into core regions could have deleterious effects on plasmid fitness (95,96). Plasmid pQBR53 had Tn4652 insert directly into genes putatively involved in horizontal transmission, and the fact we were unable to mobilise it suggests it became incapacitated as a consequence. Pre-existing TEs present in sequences can mitigate this risk by acting as a ‘safe’ target — a TE inserting within another is not disrupting an essential gene — and indeed, in 2 cases (Group IV) we identified nested transposons. This is often the case in eukaryotes (97,98). TE acquisition may not always be beneficial, as insertion into core regions could have deleterious effects on plasmid fitness (95,96). Plasmid pQBR53 had Tn4652 insert directly into genes putatively involved in horizontal transmission, and the fact we were unable to mobilise it suggests it became incapacitated as a consequence. Pre-existing TEs present in sequences can mitigate this risk by acting as a ‘safe’ target, a TE inserting within another is not disrupting an essential gene. This nesting tactic is utilised by TEs in eukaryotes (97,98) and indeed, in 2 cases (Group IV) we identified nested transposons with the pQBR collection. The activity and additional genetic material carried by TEs might also act as a burden on plasmids. Eukaryotes have defence systems that suppress activity of TEs (99), and in bacteria, H-NS proteins can act to direct transposons away from essential genes (100). A H-NS homologue was identified on pQBR26, which may act similarly to protect the plasmid from disruptive transposon activity.

Groups within the pQBR collection showed varying prevalence of defence systems and toxin-antitoxin systems, indicating divergent strategies for plasmid persistence. Erebus, a phage defence system with currently unclear mechanism of action, was uniquely present on pQBR49 amid a ∼37-kb region which was likely mobilised onto the plasmid from elsewhere, pointing to the dynamic nature of defence system reassortment on plasmids. The Group II plasmid pQBR26 possessed a CRISPR-Cas system, with spacers matching other (non-pQBR) plasmids, while the Group III plasmids all possessed a vcrx091-093 anti-CRISPR system (101), consistent with an emerging role for genome defence in plasmid competition (6,102,103). While smaller plasmids can rapidly acquire mutations to avoid restriction-modification systems, larger plasmids, like the pQBR collection, tend to acquire avoidance systems (104,105), indeed, most of the pQBR plasmids carry orphan methyltransferases that may act to provide protection against restriction endonucleases, with the exception of pQBR106. Defence systems can be valuable, but they can present a fitness cost, and may influence the ability of a plasmid to enter new hosts and to acquire and mobilise valuable new traits (106). For example, compared with other Group III plasmids, pQBR132 lacked a region containing a predicted MazEF-like anti-phage/toxin-antitoxin system, which may explain its increased rate of conjugation compared with pQBR55 from the same group.

Different pQBR plasmids varied widely in their ability to mobilise a chromosomal transposon, with some related plasmids showing order-of-magnitude variation in transposon mobilisation rates, despite similar conjugation rates, suggesting another factor was in play. Previously, we found that pQBR57 and pQBR103 plasmid acquisition directly upregulated the putative transposase of Tn6291 in a manner independent of plasmid fitness cost (33), and here we did not find any correlation between fitness costs and transposon mobilisation, suggesting that unknown genetic factors can influence rate of transposon acquisition, with some plasmids perhaps better at stimulating TEs than others, or copy numbers of TEs within the plasmid influencing expression of TE genes (107).

Our work provides a plasmid collection suitable for exploring the evolution and dynamics of gene acquisition by pre-AMR plasmids, and the diversity that can exist within and between genetically distinct plasmid groups from the same niche. This case study highlights the pervasive interplay between plasmids and TEs that can drive plasmid and microbial genome evolution. Further investigations may reveal factors that promote or reduce transposition, such as TE copy number (107), potentially uncovering genetic conflicts with particular transposons, as exist with plasmids and their hosts (33). Understanding insertion sites of transposons, within these plasmids, could reveal methods of plasmids mitigating deleterious insertions. Previously it has been shown that pQBR57 (Group IV) and pQBR103 (Group I) can coexist within a host (108), as we do not know the original hosts of these plasmids, it would useful to establish their host-range and compatibility across and within groups. This has begun to be explored with pQBR57, finding conjugation occurring between *Pseudomonas* spp. and 5 defence systems that may be a barrier to pQBR57 acquisition (109). Given the genetic diversity and variation in defence system presence across the groups, this collection could provide key insights into host-range dynamics and genome defence.

## Author contributions

VTO: Conceptualization, Data Curation, Formal Analysis, Investigation, Visualization, Writing — original draft, Writing — review & editing. JPJH: Conceptualization, Supervision, Project Administration, Funding Acquisition, Formal Analysis, Resources, Visualization, Writing — original draft, Writing — review & editing. EH: Conceptualization, Supervision, Funding Acquisition, Writing — review & editing. DR: Resources, Investigation, Writing — review & editing. RW: Resources, Writing — review & editing.

## Acknowledgements

The authors would like to thank the following people. Andy Lilley for providing the pQBR plasmids analysed here. Calvin Dytham for supervision in planning and problem solving throughout the PhD studentship associated with this work. Mitja Remus-Emsermann for providing pP19E3.1. Mike Brockhurst for preliminary investigations with strain SBW25-Tn6291::KmR. Liam Shaw and Stephen Cornell for feedback and suggestions on this work as part of the PhD viva process. Lastly, we would like to thank the team at University of Liverpool ELM lab for their support.

## Data summary

The authors confirm that all supporting data, code, and protocols have been provided within the article, through supplementary data files or GitHub repository. Raw sequencing data for all sequenced isolates have been deposited in the European Nucleotide Archive under project PRJEB71352. Accession numbers are provided in Supplementary Table S6.

## References

1. Fraikin N, Couturier A, Lesterlin C. The winding journey of conjugative plasmids toward a novel host cell. Current Opinion in Microbiology. 2024 Apr 1;78:102449. doi:10.1016/j.mib.2024.102449

2. Tokuda M, Shintani M. Microbial evolution through horizontal gene transfer by mobile genetic elements. Microbial Biotechnology. 2024;17(1):e14408. doi:10.1111/1751-7915.14408

3. Lang AS, Buchan A, Burrus V. Interactions and evolutionary relationships among bacterial mobile genetic elements. Nat Rev Microbiol. 2025 Jul;23(7):423–38. doi:10.1038/s41579-025-01157-y

4. Sen D, Van der Auwera GA, Rogers LM, Thomas CM, Brown CJ, Top EM. Broad-Host-Range Plasmids from Agricultural Soils Have IncP-1 Backbones with Diverse Accessory Genes. Applied and Environmental Microbiology. 2011 Nov 15;77(22):7975–83. doi:10.1128/AEM.05439-11

5. Rocha EPC, Bikard D. Microbial defenses against mobile genetic elements and viruses: Who defends whom from what? PLoS Biol. 2022 Jan 13;20(1):e3001514. doi:10.1371/journal.pbio.3001514

6. Samuel B, Mittelman K, Croitoru SY, Ben Haim M, Burstein D. Diverse anti-defence systems are encoded in the leading region of plasmids. Nature. 2024 Nov;635(8037):186–92. doi:10.1038/s41586-024-07994-w

7. Siedentop B, Rüegg D, Bonhoeffer S, Chabas H. My host’s enemy is my enemy: plasmids carrying CRISPR-Cas as a defence against phages. Proc Biol Sci. 2024 Jan 31;291(2015):20232449. doi:10.1098/rspb.2023.2449 PubMed PMID: 38262608; PubMed Central PMCID: PMC10805597.

8. Abbas A, Barkhouse A, Hackenberger D, Wright GD. Antibiotic resistance: A key microbial survival mechanism that threatens public health. Cell Host & Microbe. 2024 Jun 12;32(6):837–51. doi:10.1016/j.chom.2024.05.015 PubMed PMID: 38870900.

9. Acman M, Wang R, van Dorp L, Shaw LP, Wang Q, Luhmann N, et al. Role of mobile genetic elements in the global dissemination of the carbapenem resistance gene blaNDM. Nat Commun. 2022 Mar 3;13(1):1. doi:10.1038/s41467-022-28819-2

10. Cazares A, Moore MP, Hall JPJ, Wright LL, Grimes M, Emond-Rhéault JG, et al. A megaplasmid family driving dissemination of multidrug resistance in Pseudomonas. Nat Commun. 2020 Mar 13;11(1):1370. doi:10.1038/s41467-020-15081-7

11. Cyriaque V, Jacquiod S, Riber L, Abu Al-soud W, Gillan DC, Sørensen SJ, et al. Selection and propagation of IncP conjugative plasmids following long-term anthropogenic metal pollution in river sediments. Journal of Hazardous Materials. 2020 Jan 15;382:121173. doi:10.1016/j.jhazmat.2019.121173

12. Marquiegui-Alvaro A, Kottara A, Chacón M, Cliffe L, Brockhurst M, Dixon N. Genetic Bioaugmentation-Mediated Bioremediation of Terephthalate in Soil Microcosms Using an Engineered Environmental Plasmid. Microb Biotechnol. 2025 Jan;18(1):e70071. doi:10.1111/1751-7915.70071 PubMed PMID: 39801293; PubMed Central PMCID: PMC11725763.

13. Botelho J, Grosso F, Peixe L. Antibiotic resistance in Pseudomonas aeruginosa – Mechanisms, epidemiology and evolution. Drug Resistance Updates. 2019 May 1;44:100640. doi:10.1016/j.drup.2019.07.002

14. Mathers AJ, Li TJX, He Q, Narendra S, Stoesser N, Eyre DW, et al. Developing a framework for tracking antimicrobial resistance gene movement in a persistent environmental reservoir. npj Antimicrob Resist. 2024 Dec 30;2(1):50. doi:10.1038/s44259-024-00069-w

15. Sheppard AE, Stoesser N, Wilson DJ, Sebra R, Kasarskis A, Anson LW, et al. Nested Russian Doll-Like Genetic Mobility Drives Rapid Dissemination of the Carbapenem Resistance Gene blaKPC. Antimicrobial Agents and Chemotherapy. 2016 Jun;60(6):3767–78. doi:10.1128/AAC.00464-16

16. San Millan A. The journey of bacterial genes. Nat Ecol Evol. 2022 Mar 28;1–2. doi:10.1038/s41559-022-01713-2

17. Szczepanowski R, Eikmeyer F, Harfmann J, Blom J, Rogers LM, Top EM, et al. Sequencing and comparative analysis of IncP-1α antibiotic resistance plasmids reveal a highly conserved backbone and differences within accessory regions. Journal of Biotechnology. 2011 Aug 20;New Frontiers in Microbial Genome Research155(1):95–103. doi:10.1016/j.jbiotec.2010.11.018

18. Hall JPJ, Williams D, Paterson S, Harrison E, Brockhurst MA. Positive selection inhibits gene mobilization and transfer in soil bacterial communities. Nat Ecol Evol. 2017 Sep;1(9):1348–53. doi:10.1038/s41559-017-0250-3

19. Alonso-del Valle A, Toribio-Celestino L, Quirant A, Pi CT, DelaFuente J, Canton R, et al. Antimicrobial resistance level and conjugation permissiveness shape plasmid distribution in clinical enterobacteria. Proceedings of the National Academy of Sciences. 2023 Dec 19;120(51):e2314135120. doi:10.1073/pnas.2314135120

20. San Millan A, Escudero JA, Gifford DR, Mazel D, MacLean RC. Multicopy plasmids potentiate the evolution of antibiotic resistance in bacteria. Nat Ecol Evol. 2016 Nov 7;1(1):10. doi:10.1038/s41559-016-0010 PubMed PMID: 28812563.

21. Vogwill T, MacLean RC. The genetic basis of the fitness costs of antimicrobial resistance: a meta-analysis approach. Evolutionary Applications. 2015;8(3):284–95. doi:10.1111/eva.12202

22. Cazares A, Figueroa W, Cazares D, Lima L, Turnbull JD, McGregor H, et al. Pre- and postantibiotic epoch: The historical spread of antimicrobial resistance. Science. 2025 Sep 25;0(0):eadr1522. doi:10.1126/science.adr1522

23. Lilley AK, Bailey MJ, Day MJ, Fry JC. Diversity of mercury resistance plasmids obtained by exogenous isolation from the bacteria of sugar beet in three successive years. FEMS Microbiology Ecology. 1996 Aug 1;20(4):211–27. doi:10.1111/j.1574-6941.1996.tb00320.x

24. Lilley AK, Fry JC, Day MJ, Bailey MJ. In situ transfer of an exogenously isolated plasmid between Pseudomonas spp. in sugar beet rhizosphere. Microbiology. 1994;140(1):27–33. doi:10.1099/13500872-140-1-27

25. Lilley AK, Bailey MJ. The acquisition of indigenous plasmids by a genetically marked pseudomonad population colonizing the sugar beet phytosphere is related to local environmental conditions. Appl Environ Microbiol. 1997 Apr;63(4):1577–83. doi:10.1128/aem.63.4.1577-1583.1997 PubMed PMID: 16535580; PubMed Central PMCID: PMC1389558.

26. McClure NC, Weightman AJ, Fry JC. Survival of Pseudomonas putida UWC1 containing cloned catabolic genes in a model activated-sludge unit. Applied and Environmental Microbiology. 1989 Oct;55(10):2627–34. doi:10.1128/aem.55.10.2627-2634.1989

27. Bailey MJ, Lilley AK, Thompson IP, Rainey PB, Ellis RJ. Site directed chromosomal marking of a fluorescent pseudomonad isolated from the phytosphere of sugar beet; stability and potential for marker gene transfer. Molecular Ecology. 1995;4(6):755–64. doi:10.1111/j.1365-294X.1995.tb00276.x

28. Lilley AK, Bailey MJ. The transfer dynamics of Pseudomonas sp. plasmid pQBR11 in biofilms. FEMS Microbiology Ecology. 2002 Nov 1;42(2):243–50. doi:10.1111/j.1574-6941.2002.tb01014.x

29. Lilley AK, Bailey MJ. Impact of Plasmid pQBR103 Acquisition and Carriage on the Phytosphere Fitness of Pseudomonas fluorescens SBW25: Burden and Benefit. Applied and Environmental Microbiology. 1997 Apr. Located at: world. doi:10.1128/aem.63.4.1584-1587.1997

30. Hall JPJ, Wood AJ, Harrison E, Brockhurst MA. Source–sink plasmid transfer dynamics maintain gene mobility in soil bacterial communities. Proc Natl Acad Sci U S A. 2016 Jul 19;113(29):8260–5. doi:10.1073/pnas.1600974113 PubMed PMID: 27385827; PubMed Central PMCID: PMC4961173.

31. Slater FR, Bruce KD, Ellis RJ, Lilley AK, Turner SL. Heterogeneous Selection in a Spatially Structured Environment Affects Fitness Tradeoffs of Plasmid Carriage in Pseudomonads. Applied and Environmental Microbiology. 2008 May 15;74(10):3189–97. doi:10.1128/AEM.02383-07

32. Stevenson C, Hall JPJ, Brockhurst MA, Harrison E. Plasmid stability is enhanced by higher-frequency pulses of positive selection. Proceedings of the Royal Society B: Biological Sciences. 2018 Jan 10;285(1870):20172497. doi:10.1098/rspb.2017.2497

33. Hall JPJ, Wright RCT, Harrison E, Muddiman KJ, Wood AJ, Paterson S, et al. Plasmid fitness costs are caused by specific genetic conflicts enabling resolution by compensatory mutation. de Visser JAGM, editor. PLoS Biol. 2021 Oct 13;19(10):e3001225. doi:10.1371/journal.pbio.3001225

34. Hall JPJ, Wright RCT, Guymer D, Harrison E, Brockhurst MA. Extremely fast amelioration of plasmid fitness costs by multiple functionally diverse pathways. Microbiology. 2020;166(1):56–62. doi:10.1099/mic.0.000862

35. Harrison E, Guymer D, Spiers AJ, Paterson S, Brockhurst MA. Parallel Compensatory Evolution Stabilizes Plasmids across the Parasitism-Mutualism Continuum. Current Biology. 2015 Aug 3;25(15):2034–9. doi:10.1016/j.cub.2015.06.024

36. Wright RCT, Wood AJ, Bottery MJ, Muddiman KJ, Paterson S, Harrison E, et al. A chromosomal mutation is superior to a plasmid-encoded mutation for plasmid fitness cost compensation. PLOS Biology. 2024 Dec 2;22(12):e3002926. doi:10.1371/journal.pbio.3002926

37. Tett A, Spiers AJ, Crossman LC, Ager D, Ciric L, Dow JM, et al. Sequence-based analysis of pQBR103; a representative of a unique, transfer-proficient mega plasmid resident in the microbial community of sugar beet. ISME J. 2007 Aug;1(4):4. doi:10.1038/ismej.2007.47

38. Hall JPJ, Harrison E, Lilley AK, Paterson S, Spiers AJ, Brockhurst MA. Environmentally co-occurring mercury resistance plasmids are genetically and phenotypically diverse and confer variable context-dependent fitness effects. Environ Microbiol. 2015 Dec;17(12):5008–22. doi:10.1111/1462-2920.12901 PubMed PMID: 25969927; PubMed Central PMCID: PMC4989453.

39. Zhang XX, Lilley AK, Bailey MJ, Rainey PB. The indigenous Pseudomonas plasmid pQBR103 encodes plant-inducible genes, including three putative helicases. FEMS Microbiology Ecology. 2004;51(1):9–17. doi:10.1016/j.femsec.2004.07.006

40. Turner SL, Lilley AK, Bailey MJ. Two dnaB genes are associated with the origin of replication of pQBR55, an exogenously isolated plasmid from the rhizosphere of sugar beet. FEMS Microbiology Ecology. 2002 Nov 1;42(2):209–15. doi:10.1111/j.1574-6941.2002.tb01010.x

41. Viegas CA, Lilley AK, Bruce K, Bailey MJ. Description of a novel plasmid replicative origin from a genetically distinct family of conjugative plasmids associated with phytosphere microflora. FEMS Microbiol Lett. 1997 Apr 1;149(1):121–7. doi:10.1111/j.1574-6968.1997.tb10318.x

42. Hmelo LR, Borlee BR, Almblad H, Love ME, Randall TE, Tseng BS, et al. Precision-engineering the Pseudomonas aeruginosa genome with two-step allelic exchange. Nat Protoc. 2015 Nov;10(11):1820–41. doi:10.1038/nprot.2015.115

43. Schmid M, Frei D, Patrignani A, Schlapbach R, Frey JE, Remus-Emsermann MNP, et al. Pushing the limits of de novo genome assembly for complex prokaryotic genomes harboring very long, near identical repeats. Nucleic Acids Research. 2018 Sep 28;46(17):8953–65. doi:10.1093/nar/gky726

44. Li H, Durbin R. Fast and accurate long-read alignment with Burrows-Wheeler transform. Bioinformatics. 2010 Mar 1;26(5):589–95. doi:10.1093/bioinformatics/btp698 PubMed PMID: 20080505; PubMed Central PMCID: PMC2828108.

45. Danecek P, Bonfield JK, Liddle J, Marshall J, Ohan V, Pollard MO, et al. Twelve years of SAMtools and BCFtools. Gigascience. 2021 Feb 16;10(2):giab008. doi:10.1093/gigascience/giab008 PubMed PMID: 33590861; PubMed Central PMCID: PMC7931819.

46. Bankevich A, Nurk S, Antipov D, Gurevich AA, Dvorkin M, Kulikov AS, et al. SPAdes: A New Genome Assembly Algorithm and Its Applications to Single-Cell Sequencing. J Comput Biol. 2012 May;19(5):455–77. doi:10.1089/cmb.2012.0021 PubMed PMID: 22506599; PubMed Central PMCID: PMC3342519.

47. Wick RR, Schultz MB, Zobel J, Holt KE. Bandage: interactive visualization of de novo genome assemblies. Bioinformatics. 2015 Oct 15;31(20):3350–2. doi:10.1093/bioinformatics/btv383

48. Wick RR, Holt KE. Polypolish: Short-read polishing of long-read bacterial genome assemblies. PLOS Computational Biology. 2022 Jan 24;18(1):e1009802. doi:10.1371/journal.pcbi.1009802

49. Bouras G, Judd LM, Edwards RA, Vreugde S, Stinear TP, Wick RR. How low can you go? Short-read polishing of Oxford Nanopore bacterial genome assemblies. Microbial Genomics. 2024;10(6):001254. doi:10.1099/mgen.0.001254

50. Nishimura Y, Watai H, Honda T, Mihara T, Omae K, Roux S, et al. Environmental Viral Genomes Shed New Light on Virus-Host Interactions in the Ocean. mSphere. 2017;2(2):e00359–16. doi:10.1128/mSphere.00359-16 PubMed PMID: 28261669; PubMed Central PMCID: PMC5332604.

51. Schwengers O, Jelonek L, Dieckmann MA, Beyvers S, Blom J, Goesmann A. Bakta: rapid and standardized annotation of bacterial genomes via alignment-free sequence identification. Microbial Genomics. 2021;7(11):000685. doi:10.1099/mgen.0.000685

52. R Core Team. R: A Language and Environment for Statistical Computing. [Internet]. Vienna: R Foundation for Statistical Computing; 2021. Available from: https://www.R-project.org

53. Wickham H, Averick M, Bryan J, Chang W, McGowan LD, François R, et al. Welcome to the Tidyverse. Journal of Open Source Software. 2019 Nov 21;4(43):1686. doi:10.21105/joss.01686

54. Xie Y, Sarma A, Vogt A, Andrew A, Zvoleff A, Al-Zubaidi A, et al. knitr: A General-Purpose Package for Dynamic Report Generation in R [Internet]. 2024. Available from: https://cran.r-project.org/web/packages/knitr/index.html

55. Ondov BD, Treangen TJ, Melsted P, Mallonee AB, Bergman NH, Koren S, et al. Mash: fast genome and metagenome distance estimation using MinHash. Genome Biology. 2016 Jun 20;17(1):132. doi:10.1186/s13059-016-0997-x

56. Camacho C, Coulouris G, Avagyan V, Ma N, Papadopoulos J, Bealer K, et al. BLAST+: architecture and applications. BMC Bioinformatics. 2009 Dec 15;10(1):421. doi:10.1186/1471-2105-10-421

57. Sullivan MJ, Petty NK, Beatson SA. Easyfig: a genome comparison visualizer. Bioinformatics. 2011 Apr 1;27(7):1009–10. doi:10.1093/bioinformatics/btr039 PubMed PMID: 21278367; PubMed Central PMCID: PMC3065679.

58. Wishart DS, Han S, Saha S, Oler E, Peters H, Grant JR, et al. PHASTEST: faster than PHASTER, better than PHAST. Nucleic Acids Research. 2023 Jul 5;51(W1):W443–50. doi:10.1093/nar/gkad382

59. Guan J, Chen Y, Goh YX, Wang M, Tai C, Deng Z, et al. TADB 3.0: an updated database of bacterial toxin–antitoxin loci and associated mobile genetic elements. Nucleic Acids Research. 2024 Jan 5;52(D1):D784–90. doi:10.1093/nar/gkad962

60. Tesson F, Planel R, Egorov AA, Georjon H, Vaysset H, Brancotte B, et al. A Comprehensive Resource for Exploring Antiphage Defense: DefenseFinder Webservice,Wiki and Databases. Peer Community Journal. 2024 Sep 25;4. doi:10.24072/pcjournal.470

61. Tesson F, Hervé A, Mordret E, Touchon M, d’Humières C, Cury J, et al. Systematic and quantitative view of the antiviral arsenal of prokaryotes. Nat Commun. 2022 May 10;13(1):2561. doi:10.1038/s41467-022-30269-9

62. Feldgarden M, Brover V, Gonzalez-Escalona N, Frye JG, Haendiges J, Haft DH, et al. AMRFinderPlus and the Reference Gene Catalog facilitate examination of the genomic links among antimicrobial resistance, stress response, and virulence. Sci Rep. 2021 Jun 16;11(1):12728. doi:10.1038/s41598-021-91456-0 PubMed PMID: 34135355; PubMed Central PMCID: PMC8208984.

63. Ross K, Varani AM, Snesrud E, Huang H, Alvarenga DO, Zhang J, et al. TnCentral: a Prokaryotic Transposable Element Database and Web Portal for Transposon Analysis. mBio. 2021 Sep 14;12(5):10.1128/mbio.02060-21. doi:10.1128/mbio.02060-21

64. Galata V, Fehlmann T, Backes C, Keller A. PLSDB: a resource of complete bacterial plasmids. Nucleic Acids Res. 2019 Jan 8;47(D1):D195–202. doi:10.1093/nar/gky1050

65. Winsor GL, Griffiths EJ, Lo R, Dhillon BK, Shay JA, Brinkman FSL. Enhanced annotations and features for comparing thousands of Pseudomonas genomes in the Pseudomonas genome database. Nucleic Acids Res. 2016 Jan 4;44(D1):D646-653. doi:10.1093/nar/gkv1227 PubMed PMID: 26578582; PubMed Central PMCID: PMC4702867.

66. Bayliss SC, Thorpe HA, Coyle NM, Sheppard SK, Feil EJ. PIRATE: A fast and scalable pangenomics toolbox for clustering diverged orthologues in bacteria. Gigascience. 2019 Oct 1;8(10):giz119. doi:10.1093/gigascience/giz119

67. Capella-Gutiérrez S, Silla-Martínez JM, Gabaldón T. trimAl: a tool for automated alignment trimming in large-scale phylogenetic analyses. Bioinformatics. 2009 Aug 1;25(15):1972–3. doi:10.1093/bioinformatics/btp348 PubMed PMID: 19505945; PubMed Central PMCID: PMC2712344.

68. Schliep K, Paradis E, Martins L de O, Potts A, Bardel-Kahr I, White TW, et al. phangorn: Phylogenetic Reconstruction and Analysis [Internet]. 2024. Available from: https://cran.r-project.org/web/packages/phangorn/index.html

69. Thompson H. ggtreebar: Make Treemap Bar Charts with ‘ggplot2’ [Internet]. 2025. Available from: https://cran.r-project.org/web/packages/ggtreebar/index.html

70. Wong TKF, Ly-Trong N, Ren H, Baños H, Roger AJ, Susko E, et al. IQ-TREE 3: Phylogenomic Inference Software using Complex Evolutionary Models [Internet]. 2025 Apr 7. Available from: https://ecoevorxiv.org/repository/view/8916/

71. Yu G, Jones B, Arendsee Z. tidytree: A Tidy Tool for Phylogenetic Tree Data Manipulation [Internet]. 2023. Available from: https://cran.r-project.org/web/packages/tidytree/index.html

72. Simonsen L, Gordon DM, Stewart FM, Levin BR. Estimating the rate of plasmid transfer: an end-point method. Journal of General Microbiology. 1990 Nov 1;136(11):2319–25. doi:10.1099/00221287-136-11-2319

73. Huisman JS, Benz F, Duxbury SJN, de Visser JAGM, Hall AR, Fischer EAJ, et al. Estimating plasmid conjugation rates: A new computational tool and a critical comparison of methods. Plasmid. 2022 May 1;121:102627. doi:10.1016/j.plasmid.2022.102627

74. Wickham H, RStudio. forcats: Tools for Working with Categorical Variables (Factors) [Internet]. 2023. Available from: https://cran.r-project.org/web/packages/forcats/index.html

75. Hothorn T, Bretz F, Westfall P, Heiberger RM, Schuetzenmeister A, Scheibe S. multcomp: Simultaneous Inference in General Parametric Models [Internet]. 2025 [cited 2025 Feb 10]. Available from: https://cran.r-project.org/web/packages/multcomp/index.html

76. Ogle DH, Doll JC, Wheeler AP. FSA: Simple Fisheries Stock Assessment Methods [Internet]. 2025. Available from: https://cran.r-project.org/web/packages/FSA/index.html

77. Wickham H. ggplot2 [Internet]. Cham: Springer International Publishing; 2016 [cited 2025 Jan 24]. (Use R!). Available from: http://link.springer.com/10.1007/978-3-319-24277-4 doi:10.1007/978-3-319-24277-4

78. Pedersen TL. patchwork: The Composer of Plots [Internet]. 2024. Available from: https://cran.r-project.org/web/packages/patchwork/index.html

79. Wilke CO, Wiernik BM. ggtext: Improved Text Rendering Support for ‘ggplot2’ [Internet]. 2022. Available from: https://cran.r-project.org/web/packages/ggtext/index.html

80. Nishimura Y, Kaneko K, Kamijo T, Isogai N, Tokuda M, Xie H, et al. A large-scale phylogeny of replication initiation proteins illuminates plasmid macroevolutionary landscape [Internet]. bioRxiv; 2025. p. 2024.09.03.610885. Available from: https://www.biorxiv.org/content/10.1101/2024.09.03.610885v3 doi:10.1101/2024.09.03.610885

81. Hall JPJ, Botelho J, Cazares A, Baltrus DA. What makes a megaplasmid? Philosophical Transactions of the Royal Society B: Biological Sciences. 2021 Nov 29;377(1842):20200472. doi:10.1098/rstb.2020.0472

82. Molano LAG, Hirsch P, Hannig M, Müller R, Keller A. The PLSDB 2025 update: enhanced annotations and improved functionality for comprehensive plasmid research. Nucleic Acids Res. 2025 Jan 6;53(D1):D189–96. doi:10.1093/nar/gkae1095 PubMed PMID: 39565221; PubMed Central PMCID: PMC11701622.

83. Petrovski S, Blackmore DW, Jackson KL, Stanisich VA. Mercury(II)-resistance transposons Tn*502* and Tn*512*, from *Pseudomonas* clinical strains, are structurally different members of the Tn*5053* family. Plasmid. 2011 Jan 1;65(1):58–64. doi:10.1016/j.plasmid.2010.08.003

84. Berg DF van den, Costa AR, Esser JQ, Stanciu I, Geissler JQ, Zoumaro-Djayoon AD, et al. Bacterial homologs of innate eukaryotic antiviral defenses with anti-phage activity highlight shared evolutionary roots of viral defenses. Cell Host & Microbe. 2024 Aug 14;32(8):1427–1443.e8. doi:10.1016/j.chom.2024.07.007 PubMed PMID: 39094584.

85. Svet L, Parijs I, Isphording S, Lories B, Marchal K, Steenackers HP. Competitive interactions facilitate resistance development against antimicrobials. Appl Environ Microbiol. 2023 Oct 31;89(10):e0115523. doi:10.1128/aem.01155-23 PubMed PMID: 37819078; PubMed Central PMCID: PMC10617502.

86. Maier C, Huptas C, von Neubeck M, Scherer S, Wenning M, Lücking G. Genetic Organization of the aprX-lipA2 Operon Affects the Proteolytic Potential of Pseudomonas Species in Milk. Front Microbiol. 2020 Jun 10;11. doi:10.3389/fmicb.2020.01190

87. Botelho J, Lood C, Partridge SR, van Noort V, Lavigne R, Grosso F, et al. Combining sequencing approaches to fully resolve a carbapenemase-encoding megaplasmid in a Pseudomonas shirazica clinical strain. Emerg Microbes Infect. 2019;8(1):1186–94. doi:10.1080/22221751.2019.1648182 PubMed PMID: 31381486; PubMed Central PMCID: PMC6713103.

88. Makarova KS, Wolf YI, Koonin EV. Comparative genomics of defense systems in archaea and bacteria. Nucleic Acids Research. 2013 Apr 1;41(8):4360–77. doi:10.1093/nar/gkt157

89. Jain C, Rodriguez-R LM, Phillippy AM, Konstantinidis KT, Aluru S. High throughput ANI analysis of 90K prokaryotic genomes reveals clear species boundaries. Nat Commun. 2018 Nov 30;9(1):5114. doi:10.1038/s41467-018-07641-9

90. Dunivin TK, Choi J, Howe A, Shade A. RefSoil+: a Reference Database for Genes and Traits of Soil Plasmids. mSystems. 2019 Feb 26;4(1). Located at: 1752 N St., N.W., Washington, DC. doi:10.1128/msystems.00349-18

91. Botelho J, Lood C, Partridge SR, van Noort V, Lavigne R, Grosso F, et al. Combining sequencing approaches to fully resolve a carbapenemase-encoding megaplasmid in a Pseudomonas shirazica clinical strain. Emerg Microbes Infect. 2019;8(1):1186–94. doi:10.1080/22221751.2019.1648182 PubMed PMID: 31381486; PubMed Central PMCID: PMC6713103.

92. de Lima-Morales D, Chaves-Moreno D, Jarek M, Vilchez-Vargas R, Jauregui R, Pieper DH. Draft Genome Sequence of Pseudomonas veronii Strain 1YdBTEX2. Genome Announcements. 2013 May 16;1(3):10.1128/genomea.00258-13. doi:10.1128/genomea.00258-13

93. Maier C, Huptas C, von Neubeck M, Scherer S, Wenning M, Lücking G. Genetic Organization of the aprX-lipA2 Operon Affects the Proteolytic Potential of Pseudomonas Species in Milk. Front Microbiol. 2020 Jun 10;11. doi:10.3389/fmicb.2020.01190

94. Haritha A, Sagar KP, Tiwari A, Kiranmayi P, Rodrigue A, Mohan PM, et al. MrdH, a Novel Metal Resistance Determinant of Pseudomonas putida KT2440, Is Flanked by Metal-Inducible Mobile Genetic Elements. J Bacteriol. 2009 Oct;191(19):5976–87. doi:10.1128/JB.00465-09 PubMed PMID: 19648243; PubMed Central PMCID: PMC2747888.

95. He S, Hickman AB, Varani AM, Siguier P, Chandler M, Dekker JP, et al. Insertion Sequence IS26 Reorganizes Plasmids in Clinically Isolated Multidrug-Resistant Bacteria by Replicative Transposition. mBio. 2015 Jun 9;6(3):e00762–15. doi:10.1128/mBio.00762-15 PubMed PMID: 26060276; PubMed Central PMCID: PMC4471558.

96. Heieck K, Brück T. Localization of Insertion Sequences in Plasmids for L-Cysteine Production in E. coli. Genes (Basel). 2023 Jun 22;14(7):1317. doi:10.3390/genes14071317 PubMed PMID: 37510222; PubMed Central PMCID: PMC10379815.

97. Jedlicka P, Lexa M, Vanat I, Hobza R, Kejnovsky E. Nested plant LTR retrotransposons target specific regions of other elements, while all LTR retrotransposons often target palindromes and nucleosome-occupied regions: in silico study. Mobile DNA. 2019 Dec 14;10(1):50. doi:10.1186/s13100-019-0186-z

98. Levy A, Schwartz S, Ast G. Large-scale discovery of insertion hotspots and preferential integration sites of human transposed elements. Nucleic Acids Res. 2010 Mar 1;38(5):1515–30. doi:10.1093/nar/gkp1134

99. Iwasaki YW, Shoji K, Nakagwa S, Miyoshi T, Tomari Y. Transposon–host arms race: a saga of genome evolution. Trends in Genetics. 2025 Feb 19;41(5):369–89. doi:10.1016/j.tig.2025.01.009 PubMed PMID: 39979178.

100. Cooper C, Legood S, Wheat RL, Forrest D, Sharma P, Haycocks JRJ, et al. H-NS is a bacterial transposon capture protein. Nat Commun. 2024 Aug 20;15(1):7137. doi:10.1038/s41467-024-51407-5

101. Roy D, Huguet KT, Grenier F, Burrus V. IncC conjugative plasmids and SXT/R391 elements repair double-strand breaks caused by CRISPR–Cas during conjugation. Nucleic Acids Res. 2020 Jun 18;48(16):8815–27. doi:10.1093/nar/gkaa518 PubMed PMID: 32556263; PubMed Central PMCID: PMC7498323.

102. Zheng H, Payne L, He W, Mestre MR, Yang L, Dechesne A, et al. Plasmids as persistent genetic reservoirs of bacterial defense systems in wastewater treatment. Microbiome. 2026 Jan 26;14(1):50. doi:10.1186/s40168-025-02297-2

103. Sünderhauf D, Ringger JR, Payne LJ, Pinilla-Redondo R, Gaze WH, Brown SP, et al. CRISPR-Cas is beneficial in plasmid competition, but limited by competitor toxin–antitoxin activity when horizontally transferred. PLOS Biology. 2026 Feb 19;24(2):e3003658. doi:10.1371/journal.pbio.3003658

104. Shaw LP, Rocha EPC, MacLean RC. Restriction-modification systems have shaped the evolution and distribution of plasmids across bacteria. Nucleic Acids Research. 2023 Jul 21;51(13):6806–18. doi:10.1093/nar/gkad452

105. Tesson F, Huiting E, Wei L, Ren J, Johnson M, Planel R, et al. Exploring the diversity of anti-defense systems across prokaryotes, phages and mobile genetic elements. Nucleic Acids Research. 2025 Jan 13;53(1):gkae1171. doi:10.1093/nar/gkae1171

106. Zaayman M, Wheatley RM. Fitness costs of CRISPR-Cas systems in bacteria. Microbiology. 2022;168(7):001209. doi:10.1099/mic.0.001209

107. Sastre-Dominguez J, DelaFuente J, Toribio-Celestino L, Herencias C, Herrador-Gómez P, Costas C, et al. Plasmid-encoded insertion sequences promote rapid adaptation in clinical enterobacteria. Nat Ecol Evol. 2024 Nov;8(11):2097–112. doi:10.1038/s41559-024-02523-4

108. Kottara A, Carrilero L, Harrison E, Hall JPJ, Brockhurst MA. The dilution effect limits plasmid horizontal transmission in multispecies bacterial communities. Microbiology. 2021;167(9):001086. doi:10.1099/mic.0.001086

109. 109. Marquiegui-Alvaro A, Kottara A, Thomas MJN, Scarampi A, Chacón M, Brockhurst M, et al. Using auxotrophic donor strains to explore pQBR57 plasmid host range among environmental soil bacterial isolates [Internet]. bioRxiv; 2026. p. 2026.02.11.702040. Available from: https://www.biorxiv.org/content/10.64898/2026.02.11.702040v1 doi:10.64898/2026.02.11.702040

