## Supplementary Text for "The pQBR mercury resistance plasmids: a model set of sympatric environmental mobile genetic elements"

### *Supplementary Methods*

#### ***Creation of P. fluorescens SBW25 strain contain kanamycin resistant marker on Tn6291***

The suicide vector pTS-1_Tn6291::KmR was generated from pTS-1 (1) by amplifying 900 bp upstream and downstream fragments homologous to Tn6291 from *P. fluorescens* SBW25, a 1219 bp fragment containing the *aphA1* gene for kanamycin phosphotransferase and associated promoter region from pUCP18-KmR (2), and the lambda t0 and rrnB T1 terminators from pUC18T-mini-Tn7T-Gm (3). Fragments contained appropriate overhangs for combination into the BamHI/HindIII site of pTS-1 by HiFi assembly (New England Biolabs). Clones were verified by PCR and Sanger sequencing, and the plasmid introduced into *P. fluorescens* SBW25 by electroporation (4). Single crossovers were selected on plates supplemented with tetracycline (100 µg/ml) and tested by colony PCR, before growing and spreading samples on plates supplemented with kanamycin and 10% w/v sucrose to select for second crossovers. The resulting candidate clones were tested for kanamycin resistance, tetracycline susceptibility, and the production of appropriately-sized bands by PCR. The final construct was validated by whole genome sequencing (Illumina 2x250 bp) and found to have incorporated the *aphA1* gene with corresponding promoter and terminator as expected at position 2076740 in the reference sequence AM181176. This location was selected as being downstream of open reading frames on the forward and reverse strands (PFLU_1901, a putative cytochrome D oxidase subunit II, and PFLU_1902, a putative inorganic pyrophosphatase) to minimise potential polar effects. No second-site mutations were detected.

#### ***Sequencing and assembly of pQBR plasmids***

We used a hybrid method to produce plasmid sequences for subsequent analysis. Whole genome SPAdes assemblies of Illumina reads, viewed in Bandage (0.9.0), resulted in closed contigs for the Group I plasmid pQBR103, the Group IV plasmid pQBR102, and the Group III plasmids pQBR55, pQBR132 and pQBR28. The Group III plasmid pQBR53 could also be resolved from the Illumina assembly owing to the presence of a plasmid backbone with >99.99% nucleotide identity to the known Group III plasmid pQBR55, forming junctions with a contig matching the chromosomal transposon Tn4652 (also present, but in a different location, in pQBR55). The path of the plasmid backbone and transposon contigs was considered a closed assembly for pQBR53.

ONT sequencing was performed on the following pools (Groups are provided in brackets): pQBR30 (IV), pQBR5 (I) and pQBR24 (II); pQBR57 (IV), pQBR4 (I) and pQBR23

(II); pQBR124 (I), pQBR56 (IV) and pQBR105 (NA); pQBR49 (I) and pQBR150 (IV). Plasmids pQBR51 (IV), pQBR11 (I), pQBR47 (I), pQBR106 (I), pQBR127 (IV), pQBR50 (I), pQBR43 (I), and pQBR26 (II) were each sequenced individually. ONT assemblies, produced by Flye and polished using Medaka, were further polished the corresponding Illumina reads using Polypolish 0.5.0, and direct repeats at the start and end of all assemblies were identified by ccfind (5) and removed as assembly artefacts. Overall, these approaches produced closed plasmid sequences for 20/28 UWC1(pQBR) strains. Short-read sequencing of UWC1(pQBR127) revealed replicons that both resembled Group I and Group III plasmids, but long-read sequencing showed a distinct Group III replicon and no Group I replicon. We concluded that two plasmids were present in the strain used for Illumina sequencing, of which only one remained for subsequent long-read and phenotypic analysis, which we labelled pQBR127. Sequencing of UWC1(pQBR26) identified two separate plasmid sequences harbouring mercury resistance elements: one sequence had coverage similar to the chromosome and transferred into recipients, whereas the other, which had high identity (>99.99% nucleotide identity) to the other Group II plasmids pQBR23 and pQBR24, was at lower coverage (~0.75x chromosomal coverage) and was not detected to transfer (PCR on transconjugants, n = 8 attempts), so we labelled the former plasmid pQBR26.

In one case (pQBR11), we could not identify any matching Group I contigs from long-read sequencing, but a resolved sequence could be generated from three contigs of the Illumina assembly, which were oriented with reference to pQBR103 and joined to generate the complete draft. In 4 cases (pQBR43, pQBR47, pQBR50, pQBR106), closed sequences could not be obtained, owing to the presence of long (~40 kb) repeated Tn6290 transposon sequences in the plasmid and the chromosome that could not be resolved by long-read sequencing. For pQBR47 and pQBR106, the Flye assembly of ONT reads suggested that the plasmids had integrated into the chromosome, possibly through Tn6290 recombination. The chromosomal region was removed for the subsequent plasmid analyses. For pQBR43, the single putative plasmid contig was bound by Tn6290-like sequences at either end, and for pQBR50 the assembly was in 2 contigs, each of which was bordered by Tn6290 copies. In both of these assemblies, Tn6290 coverage was higher than plasmid and chromosomal coverage, suggesting the presence of multiple Tn6290 copies in the genome. Sequences were oriented using EMBOSS to place the first base of the first codon of a putative replicase, identified by querying a BLAST database of putative replicases from previously-sequenced pQBR plasmids, at the first position on the forward strand, prior to annotation using bakta (v1.8.2, full database v.5.0.0).

In three cases (pQBR1, pQBR8, pQBR58), long deep contigs could not be obtained during the initial comparative analysis, and instead short contigs matching only a putative mercury resistance transposon, and/or longer contigs at lower coverage (<0.5x chromosome coverage) were identified. In these cases, we concluded that the plasmid was at intermediate frequency in the sequenced samples, indicative of plasmid loss from the population following capture of mercury resistance to the chromosome, and attempts to reisolate the plasmid by conjugation into a new host were unsuccessful.

Three plasmids, sequenced in an earlier analysis (6), were sequenced again from the original stock: pQBR55 (Genbank: LN713927.1), pQBR57 (Genbank: LN713926.1) and pQBR103 (AM235768.1). While the resolved sequence of pQBR103 was identical between the original and resequenced samples, pQBR57 and pQBR55 varied in the presence of transposons, with the resequenced pQBR57 harbouring Tn4652 (bases 136385-153397, similar but not identical to patterns previously observed (7) and the resequenced pQBR55 lacking Tn4652. The resequenced assemblies were labelled pQBR57R and pQBR55R. We included in our analysis the published sequence of pQBR44 (Genbank: CDLQ010000001-CDLQ010000002) (6), treating the two contigs as 'drafts', as described above.

***Measuring plasmid fitness effects***

Plasmid-containing SBW25-Tn6291::KmR strains to be tested were competed against a reference plasmid-free SBW25::SmR-*lacZ*. A control comparison between plasmid-free SBW25-Tn6291::KmR and SBW25::SmR-*lacZ* was conducted to test for any effects of the KmR marker. Single colonies were obtained from each strain and cultured in 5 ml KB broth overnight. The following day, competitions were mixed at approximately 1:1 ratio, diluted 1:100 into fresh KB broth, and incubated for 24 hours. Dilutions of the initial competition mix were plated onto KB agar containing X-gal to obtain start counts. After incubation, competitions were diluted and spread on KB agar containing X-gal to obtain end counts. Competitors could then be distinguished using colour, with SBW25::SmR-*lacZ* appearing blue compared to the white competitor strains.

Plate counts were transformed into competitive fitness value "w" using the following equations(8):

$$m = \ln\left(\frac{N/N_0}{T}\right)$$

$$w = \frac{m_{test}}{m_{reference}}$$

N is the total colonies of a strain at the end of the experiment and $N_0$ is the total colonies of strain at the start of the experiment. T is the experiment duration, in this case 24 hours.

### *Supplementary Results and Discussion*

#### *Substitutions in Tn5042 across collection and mer operons in pQBR26 and pQBR105*

All copies of Tn5042 within the pQBR collection had 35 nucleotide substitutions (Supplementary: Table S2), 7 insertions and 4 deletions compared with the reference sequence (AJ563380.2) Group I plasmid copies (except pQBR11 and pQBR49) contained an addition G296>T substitution (within *merR*) and pQBR132 contained both a C733>A (intergenic region) and G4635>T (within hypothetical protein).

Tn5042 was not present in pQBR26 and pQBR105, TnCentral could not identify any complete mercury transposons within these sequences. Plasmid pQBR26 contains a *merRTPCADE* operon as well as separate *merED, merPTR* and *merR* genes and pQBR105 encodes *merR, merRTP* and *merPTR*. These *mer* genes are all flanked by putative transposase genes suggesting they are likewise present on TEs.

#### *Details on transposon content the pQBR plasmids of Tn4652 and Tn6290*

Tn4652 harbours a metal-resistant determinant, *mrdH*, that confers cadmium and zinc resistance in *P. putida* KT2440, and *mreA*, a metal-sensing transcriptional repressor which was shown to be conserved in association with *mrdH* (9). Annotation for bakta suggests the Tn6290 sequence encodes copper resistance genes, putative copper efflux transporters as well as other heavy metal transportation genes. Tn6291 contains many hypothetical proteins but also a putative *prtN* gene, which in *Pseudomonas aeruginosa* is involved in activate of pyocin production (10). Besides the full-length TEs described in the main text, by comparing sequences with those in TnCentral we also found some matches to variants of Tn4662 amongst the Group II plasmids, which may contain a toxin-antitoxin system (11).

#### *Gene content variations in the Group IV plasmids pQBR30 and pQBR102*

Plasmid pQBR30 lacked several ‘phage-related’ genes present in other Group IV plasmids, while pQBR102 did not encode a ferredoxin-NADP+ reductase, homologues of which are involved in redox metabolism (12). Several genes within pQBR30 and pQBR102 were truncated, including a HNH endonuclease in pQBR102 (pQBR102_01690). Unique to pQBR30 and pQBR102 is an autotransporter outer membrane beta-barrel domain-containing protein (PQBR102_01340, PQBR30_01205).

***Plasmid mobilisation of chromosomal traits varies across pQBR plasmids, and is not determined by conjugation rate – comparison of ratios***

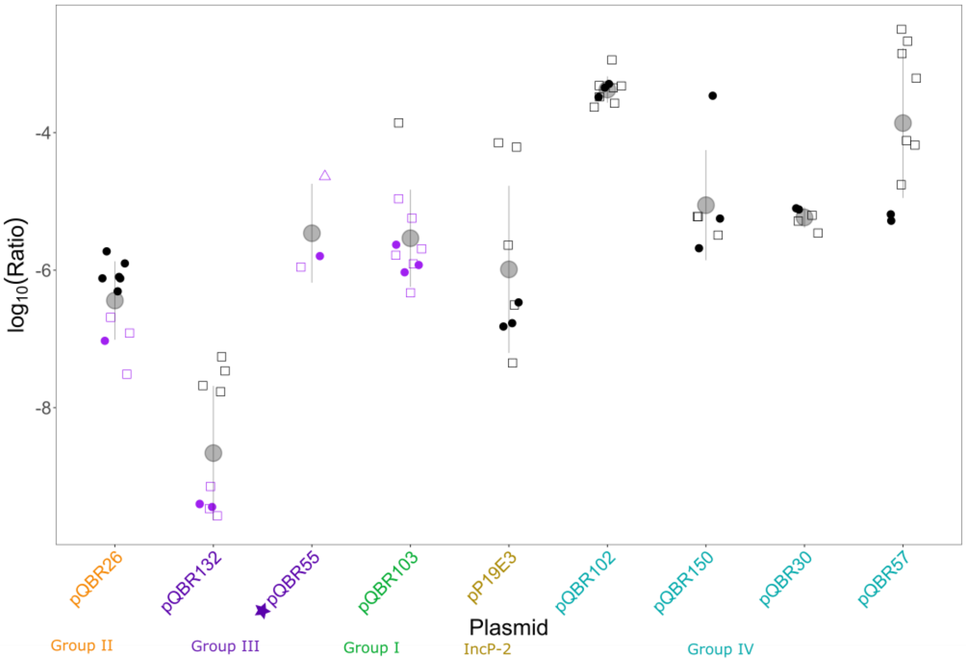


**Figure S1** Conjugation rate is not a good predictor of transposon mobilisation. Ratios calculated by dividing plasmid transposon mobilisation rate by plasmid mobilisation rate, which were used for analysis. This figure represents a summary of two experiments performed a month apart, with 2-3 different donor transconjugant strains for each plasmid identified by shape. Large points represent average measurement. Purple points indicate where transposon mobilisation measurement was below limit of detection for experiment and given an arbitrary small value. Colours of plasmid labels represents the group the plasmid belongs to, labelled beneath. Star (pQBR55) indicates that data for this plasmid was collected separately from the rest of the dataset under similar experimental conditions.

***Comparative Fitness of pQBR plasmids***

To investigate whether these patterns could be explained by the effect of the plasmids on cell fitness, we conducted growth curve and competitive fitness assays. We focused on the Group IV plasmids, owing to their significant within-group variation in transposon mobilisation rates.

All plasmid-carrying strains suffered a decrease in growth compared with a plasmid-free control (measured as area under the curve (AUC): ANOVA: $F_{6,35} = 28.04$, $p < 0.01$; post-hoc pairwise comparisons between plasmid-carrying strains and control all $p < 0.03$; and maximum per capita growth rate: ANOVA: $F_{6,35}=8.724$, $p<0.01$). These growth effects varied; plasmids pQBR103, pQBR102 and pQBR150 resulted in significantly reduced growth compared with some or all of the Group IV plasmids (Figure S2 A & B).

We did not detect any differences in fitness costs of the Group IV plasmids in head-to-head competitive fitness assays (Figure S2C), though fitness costs relative to a plasmid-free control were also not detected, possibly owing to high conjugation rates obscuring differences in this assay. but intergroup differences were observed, e.g. between Group IV and pQBR103 (Group I). There was no obvious correlation between plasmid fitness effects and the efficiency of the plasmid in mobilising a chromosomal locus, suggesting that the striking differences in transposon acquisition and mobilisation are due to more subtle differences in plasmid activity or gene content.

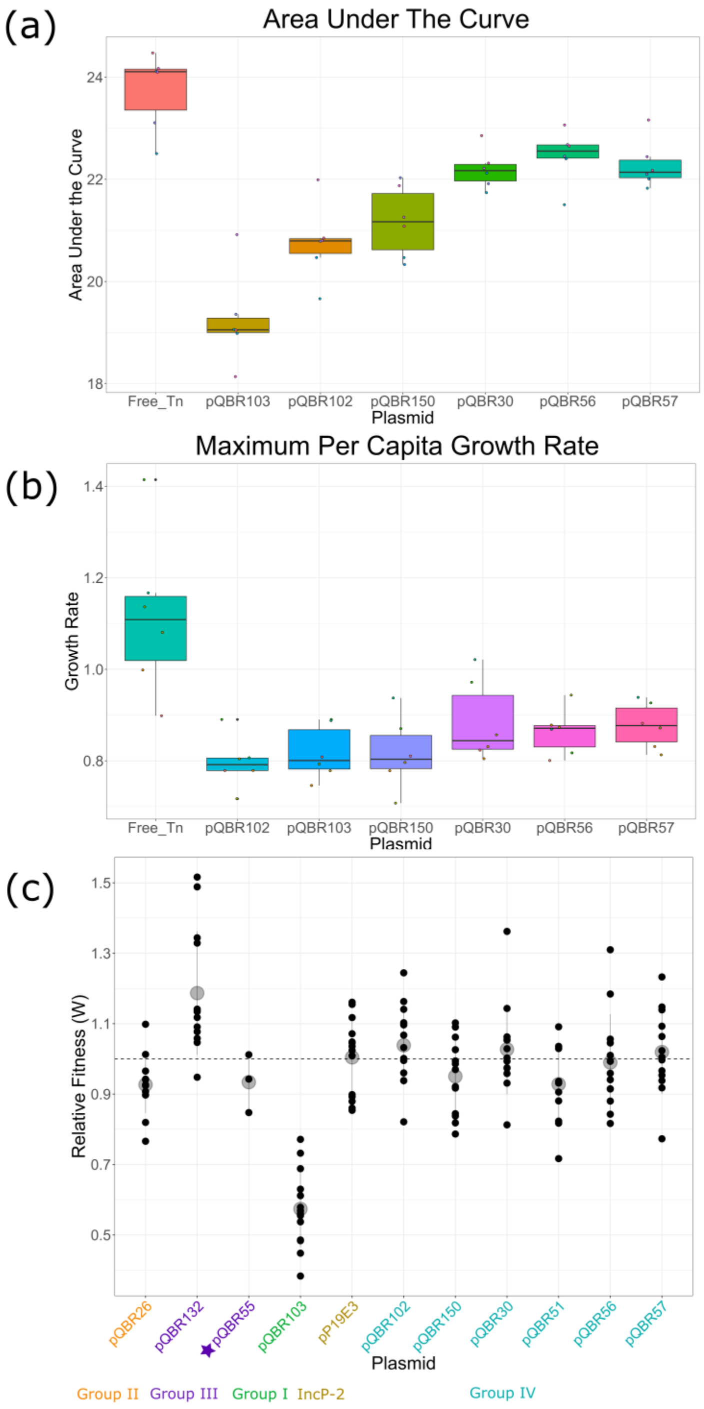


**Figure S2 A.** pQBR plasmids Group IV and pQBR103 incur a fitness cost compared to plasmid free strain **B.** Maximum per capita growth rate was reduced in plasmid carrying strains compared to a plasmid free strain. **C.** Intergroup plasmid fitness costs were greater than intra-group differences. Group IV plasmid fitness cost did not vary significantly. Colours of plasmid labels represents the group the plasmid belongs to, labelled beneath. Star (pQBR55) indicates that data for this plasmid was collected separately from the rest of the dataset under similar experimental conditions.

### References


1. Campilongo R, Fung RKY, Little RH, Grenga L, Trampari E, Pepe S, et al. One ligand, two regulators and three binding sites: How KDPG controls primary carbon metabolism in Pseudomonas. PLOS Genetics. 2017 Jun 28;13(6):e1006839. doi:10.1371/journal.pgen.1006839

2. Schweizer HP. Escherichia-Pseudomonas shuttle vectors derived from pUC18/19. Gene. 1991 Jan 2;97(1):109–21. doi:10.1016/0378-1119(91)90016-5 PubMed PMID: 1899844.

3. Choi KH, Gaynor JB, White KG, Lopez C, Bosio CM, Karkhoff-Schweizer RR, et al. A Tn7-based broad-range bacterial cloning and expression system. Nat Methods. 2005 Jun;2(6):443–8. doi:10.1038/nmeth765

4. Choi KH, Ayush K, Schweizer HP. A 10-min method for preparation of highly electrocompetent Pseudomonas aeruginosa cells: Application for DNA fragment transfer between chromosomes and plasmid transformation. Journal of Microbiological Methods. 2006 Mar 1;64(3):391–7. doi:10.1016/j.mimet.2005.06.001

5. Nishimura Y. yosuken/ccfind [Ruby] [Internet]. 2024 [cited 2025 Feb 28]. Available from: https://github.com/yosuken/ccfind

6. Hall JPJ, Harrison E, Lilley AK, Paterson S, Spiers AJ, Brockhurst MA. Environmentally co-occurring mercury resistance plasmids are genetically and phenotypically diverse and confer variable context-dependent fitness effects. Environ Microbiol. 2015 Dec;17(12):5008–22. doi:10.1111/1462-2920.12901 PubMed PMID: 25969927; PubMed Central PMCID: PMC4989453.

7. Hall JPJ, Williams D, Paterson S, Harrison E, Brockhurst MA. Positive selection inhibits gene mobilization and transfer in soil bacterial communities. Nat Ecol Evol. 2017 Sep;1(9):1348–53. doi:10.1038/s41559-017-0250-3

8. Lenski RE. Quantifying fitness and gene stability in microorganisms. Biotechnology. 1991;15:173–92. doi:10.1016/b978-0-409-90199-3.50015-2 PubMed PMID: 2009380.

9. Haritha A, Sagar KP, Tiwari A, Kiranmayi P, Rodrigue A, Mohan PM, et al. MrdH, a Novel Metal Resistance Determinant of Pseudomonas putida KT2440, Is Flanked by Metal-Inducible Mobile Genetic Elements. J Bacteriol. 2009 Oct;191(19):5976–87. doi:10.1128/JB.00465-09 PubMed PMID: 19648243; PubMed Central PMCID: PMC2747888.

10. Matsui H, Sano Y, Ishihara H, Shinomiya T. Regulation of pyocin genes in Pseudomonas aeruginosa by positive (prtN) and negative (prtR) regulatory genes. J Bacteriol. 1993 Mar 1;175(5):1257–63. doi:10.1128/jb.175.5.1257-1263.1993 PubMed PMID: 8444788; PubMed Central PMCID: PMC193209.

11. Yano H, Miyakoshi M, Ohshima K, Tabata M, Nagata Y, Hattori M, et al. Complete Nucleotide Sequence of TOL Plasmid pDK1 Provides Evidence for Evolutionary History of IncP-7 Catabolic Plasmids. J Bacteriol. 2010 Sep;192(17):4337–47. doi:10.1128/JB.00359-10 PubMed PMID: 20581207; PubMed Central PMCID: PMC2937381.

12. Monchietti P, López Rivero AS, Ceccarelli EA, Catalano-Dupuy DL. A new catalytic mechanism of bacterial ferredoxin-NADP+ reductases due to a particular NADP+ binding mode. Protein Sci. 2021 Oct;30(10):2106–20. doi:10.1002/pro.4166 PubMed PMID: 34382711; PubMed Central PMCID: PMC8442965.
